# Multiple morphogens and rapid elongation promote segmental patterning during development

**DOI:** 10.1101/2021.04.22.440966

**Authors:** Yuchi Qiu, Lianna Fung, Thomas F. Schilling, Qing Nie

**Author notes:** Corresponding authors: TF Schilling (TFS) and Q Nie (QN).

## Abstract

The vertebrate hindbrain is segmented into rhombomeres (r) initially defined by distinct domains of gene expression. Previous studies have shown that noise-induced gene regulation and cell sorting are critical for the sharpening of rhombomere boundaries, which start out rough in the forming neural plate (NP) and sharpen over time. However, the mechanisms controlling simultaneous formation of multiple rhombomeres and accuracy in their sizes are unclear. We have developed a stochastic multiscale cell-based model that explicitly incorporates dynamic morphogenetic changes (i.e. convergent-extension of the NP), multiple morphogens, and gene regulatory networks to investigate the formation of rhombomeres and their corresponding boundaries in the zebrafish hindbrain. During pattern initiation, the short-range signal, fibroblast growth factor (FGF), works together with the longer-range morphogen, retinoic acid (RA), to specify all of these boundaries and maintain accurately-sized segments with sharp boundaries. At later stages of patterning, we show a nonlinear change in the shape of rhombomeres with rapid left-right narrowing of the NP followed by slower dynamics. Rapid initial convergence improves boundary sharpness and segment size by regulating cell sorting and cell fate both independently and coordinately. Overall, multiple morphogens and tissue dynamics synergize to regulate the sizes and boundaries of multiple segments during development.

**Author Summary:** In segmental pattern formation, chemical gradients control gene expression in a concentration-dependent manner to specify distinct gene expression domains. Despite the stochasticity inherent to such biological processes, precise and accurate borders form between segmental gene expression domains. Previous work has revealed synergy between gene regulation and cell sorting in sharpening borders that are initially rough. However, it is still poorly understood how size and boundary sharpness of *multiple* segments are regulated in a tissue that changes dramatically in its morphology as the embryo develops. Here we develop a stochastic multiscale cell-base model to investigate these questions. Two novel strategies synergize to promote accurate segment formation, a combination of long- and short-range morphogens plus rapid tissue convergence, with one responsible for pattern initiation and the other enabling pattern refinement.

## INTRODUCTION

A fundamental question in developmental biology is how cell fate decisions are coordinated with tissue morphogenetic changes during pattern formation. During embryogenesis, cells must convert concentration-dependent positional information from diffusible chemical morphogens into coordinated cell fate decisions [1–3]. Mathematical models have successfully integrated tissue morphogenesis and spatial signaling during patterning of embryonic segments in both flies [4, 5] and vertebrates [6–8], in structures such as the wing imaginal discs in *Drosophila* [9–11], as well as the limb buds [12], neural tube [13–15], hindbrain [16–18], pharyngeal arches [19], skin [20, 21] and hair follicles [22] of vertebrates.

Understanding stochastic effects in patterning systems, particularly how precision is achieved in spite of biological noise in gene expression and spatial signals, is a major challenge in developmental biology. Noise attenuation mechanisms in gene expression have been widely explored in diverse cellular networks [23, 24]. For spatial signals, binding with membrane-bound non-signaling entities [25], regulation of gradient steepness by ligand shuttling [26, 27] and self-regulated ligand uptake [28, 29] can reduce spatial variation in morphogen gradients. Anti-parallel morphogens [14] and gene regulatory networks [30–32] that translate noisy spatial signals into cell fate decisions can also reduce patterning errors. Interestingly, noise in gene expression can counteract other stochastic effects (e.g. noise in morphogen levels) to improve pattern formation precision [17, 33]. In addition to these molecular strategies, pattern precision can be improved through cellular strategies, such as cell sorting driven by cell-cell interactions [16, 20] or “community effects” of signals from adjacent cells [34]. Previous modeling studies have often neglected to take into account rapid changes in tissue morphology, and how the interaction between these and noise attenuation mechanisms impacts pattern precision remains poorly understood.

The embryonic zebrafish hindbrain is a powerful model system to study the roles of gene regulation, stochasticity, cell sorting, and tissue morphogenesis in segmental pattern formation. Neurons in the hindbrain contribute to the cranial nerves that innervate the face and neck and control many involuntary functions, such as feeding and breathing. These neurons arise in early embryonic segments, called rhombomeres (r), that progressively subdivide along the anterior-posterior (A-P) axis [35]. Initial gene expression domains that specify segmental cell identities in rhombomeres 1-7 (r1-7) have rough borders that subsequently sharpen [17, 36]. Several spatial signals provide positional information for the establishment of rhombomeres, such as retinoic acid (RA) [28, 37–40] and fibroblast growth factors (FGFs) [41–45]. These signals regulate numerous transcription factors, including *hox* genes, *krox20*, *val*, *vhnf1* and *irx*, with rhombomere-specific expression domains that specify rhombomere cell identity [46–48]. Rhombomere-specific gene regulatory networks commit cells to distinct segmental fates and can switch their identities from one segment to another by interpreting RA and FGF signals [34, 49]. In addition, the complementary segmental expression of Ephrins and Eph receptors drives boundary sharpening by regulating cell sorting with differential adhesion/repulsion [50–53]. Previous computational models incorporating one morphogen, RA, and two transcription factors, *hoxb1a* and *krox20*, successfully mimic boundary sharpening in r3-5 by incorporating gene regulation and cell sorting [16, 17]. During the period when these boundaries sharpen, the hindbrain grows and elongates, often termed as convergent extension, where the hindbrain narrows along the left-right (L-R) axis and elongates along the anterior-posterior (A-P) axis [49, 54]. However, previous computational models have shown that tissue elongation disrupts the sharpening of rhombomere boundaries [17]. It remains unclear in any segmented tissue how multiple segments simultaneously form with accurate sizes and sharp boundaries during such morphogenetic tissue dynamics.

Here we consider hindbrain patterning across multiple stages, from pattern initiation to sharpening, across multiple segments (r2-6) and in the context of morphogenetic changes in hindbrain size and shape. We include a second morphogen in our model, FGF produced in r4, and two additional transcription factors, *vhnf1* and *irx3*. We find that FGF produced in r4 is critical to specify the r5/r6 boundary, and to achieve a robust five-segment pattern with accurate segment sizes and sharp boundaries despite variations in initial gene expression. At later stages of patterning, we show experimentally that L-R narrowing of the zebrafish hindbrain occurs rapidly at first (11-12 hours post fertilization (hpf)), but the narrowing rate drops rapidly over the following two hours (12-14 hpf). Interestingly, comparisons of hindbrain pattern formation in our model under different convergence rates suggest that such a rapid initial convergence facilitates robust patterning, both in the accuracy of segment size and boundary sharpness. This rapid initial convergence helps mediate a trade-off between boundary sharpness and segment size. Together, the cooperation between two morphogens and morphogenetic dynamics effectively regulates robust segmental patterning in the zebrafish hindbrain.

## RESULTS

### A stochastic multiscale cell-based model for hindbrain segmentation

To address how multiple morphogens and dynamics of tissue morphogenesis contribute to segmental pattern formation in the hindbrain, we developed a computational model that incorporates stochastic gene regulation, cell sorting and tissue shape changes (Fig 1). We first provide an overview of the elements, assumptions, and metrics included in our models (details see **Methods** and **Supplementary Materials)**.

**Fig 1.**
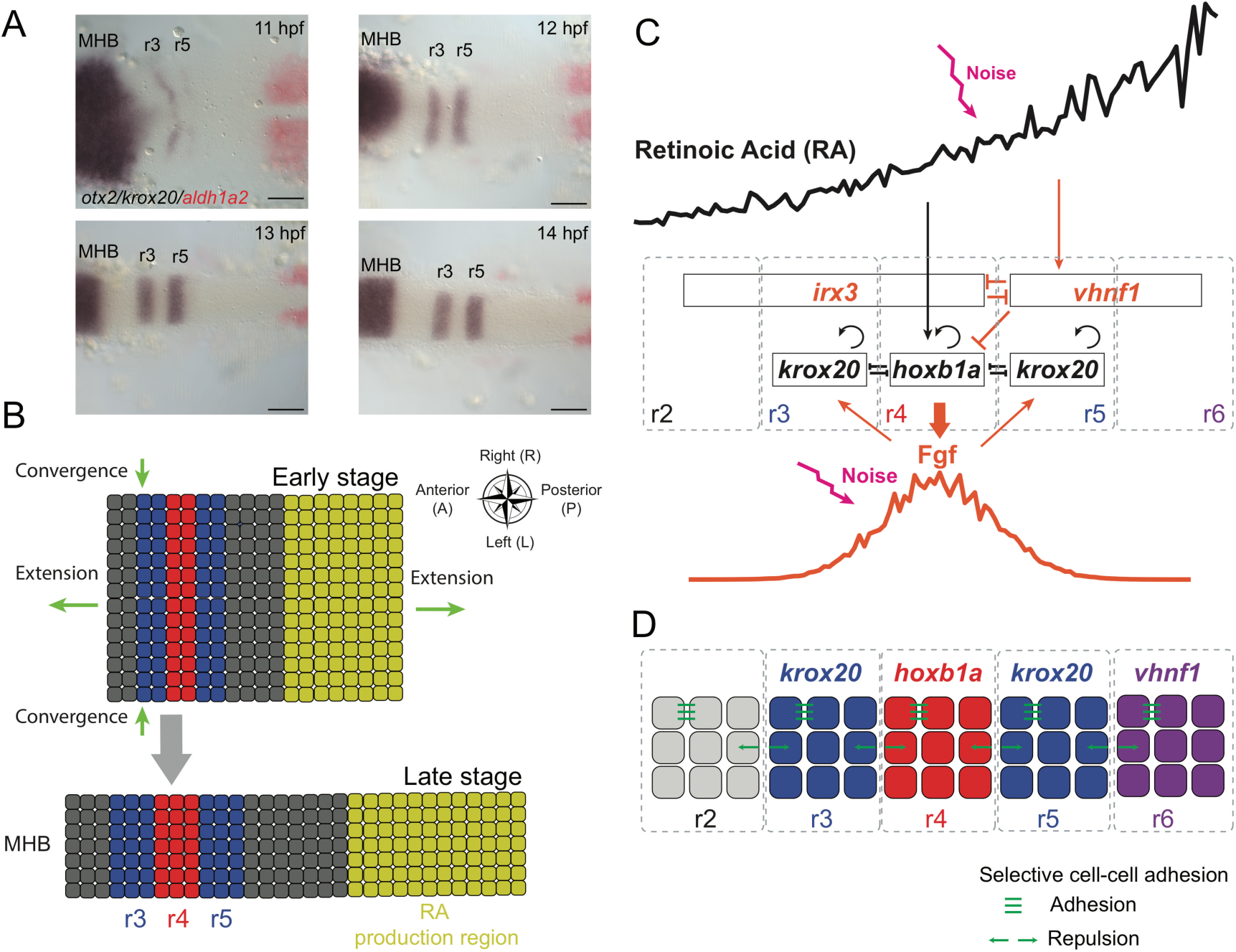
Model schematic and zebrafish hindbrain morphology. **(A)** Two-color whole mount in situ hybridization of embryonic zebrafish hindbrains for *otx2* (purple, anterior region, far left), *krox20* (purple segments, center) and *aldh1a2* (red, far right) transcripts from 11 to 14 hpf. *otx2* marks the midbrain-hindbrain boundary (MHB), *krox20* marks r3 and r5 and *aldh1a2* marks the RA production region. Embryos are flat-mounted and shown in dorsal view with anterior to the left. Scale bars: 100 µm. **(B)** Illustration depicting convergent-extension of the hindbrain. The entire rectangular region, including r1-7 and the RA production region, constitutes the morphogen domain. The hindbrain narrows in the L-R direction (width) and elongates in the A-P direction. **(C)** Gene regulatory network used to model hindbrain patterning in r2-6. Genes and morphogens with black font were previously used for modeling the r3-r5 pattern [17], while additional factors considered in this model are shown in orange font. Pointed arrows depict up-regulation/activation and blunt arrows depict down-regulation/inhibition. Two morphogens, retinoic acid (RA) and Fibroblast Growth Factor (FGF) diffuse and form two distinct gradients to govern downstream gene expression. **(D)** Illustration depicting r2-6 and distinct identities (i.e. gene expression signatures) underlying selective cell sorting. Cells in r3 and r5 (blue) express *krox20* and cells in r4 (red) express *hoxb1a*, while both *krox20* and *hoxb1a* levels are low in r2 and r6. Cells in r6 (purple) have high *vhnf1* expression. Cells with the same segmental identity attract each other and cells with different identities repulse each other.

#### Hindbrain morphogenesis and computational domains

During embryogenesis from 11-14 hpf, the zebrafish hindbrain narrows along the L-R axis and elongates along the A-P axis [49, 54]. The midbrain-hindbrain boundary (MHB) (anterior to the hindbrain and adjacent to r1) and the RA production region (posterior to the hindbrain and adjacent to r7) provide A-P landmarks for the region that forms the hindbrain (Fig 1A). To quantify these changes in size and shape, we performed whole mount in situ hybridization with markers for the MHB (*otx2*), r3 and r5 (*krox20*) and RA production (*aldh1a2*). At each hour between 11-14 hpf, we measured hindbrain width at the level of r4 as well as the A-P distance between the *otx2* and *aldh1a2* domains (Fig 2A, B). Based on these experimental measurements, we modeled the hindbrain (r1-r7) along with the RA production region as a rectangle, subsequently referred to as the morphogen domain (Fig 1B). Two morphogens, RA and FGF, were modeled in the morphogen domain. Due to the expensive computational cost, instead of modeling all cells in the morphogen domain for this study, we explicitly modeled cells in a smaller region containing r2-r6, subsequently referred to as the tissue domain.

**Fig 2.**
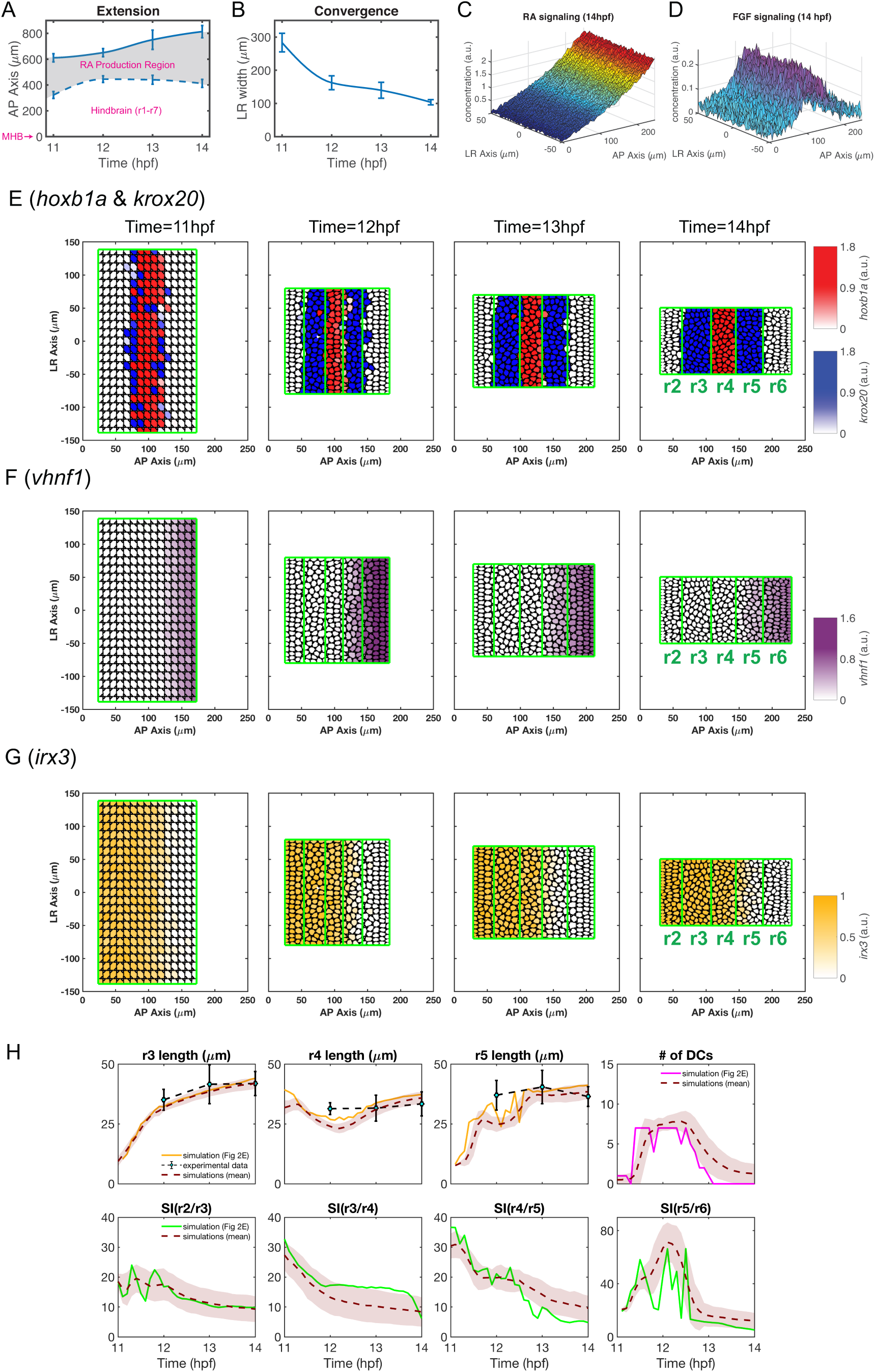
A baseline simulation mimics rhombomere boundary sharpening. **(A,B)** Experimental measurements of hindbrain dimensions along the A-P **(A)** and L-R **(B)** axes at 11, 12, 13 and 14 hpf. Error bars represent standard deviation. Cubic interpolation is used to obtain the smooth curves used in the model. **(A)** A-P hindbrain length was measured from the posterior edge of the mid-hindbrain boundary (MHB) to the anterior edge of the RA production region. A-P length of the RA production region was based on measurements of the *aldh1a2* expression domain. **(B)** L-R hindbrain width was measured at the A-P position of r4. **(C,D)** Predicted noisy distributions of morphogen signaling at 14 hpf **(C)** RA ([*RA*]*_in_*). **(D)** FGF ([*FGF*]*_signal_*). **(E-G)** Time series of gene expression in r2-6 (the hindbrain is represented as a rectangle for simplification): **(E)** *hoxb1a* (red) and *krox20* (blue), **(F)** *vhnf1* (purple), **(G)** *irx3* (yellow). **(H)** Quantifications of rhombomere length, number of dislocated cells (DCs) and sharpness indices (SIs) versus time. Rhombomere lengths (r3-5), and SIs for four boundaries (SI(r2/r3), SI(r3/r4), SI(r4/r5), SI(r5/r6)) and DC number in multiple simulations (n=100): ‘solid line’: quantities for the simulation shown in **(E)**; ‘brown dashed line’ indicates the average and the width of ‘brown shade” indicates standard deviation; ‘black dashed line’ represents rhombomere lengths from experimental measurements and the error bars represent standard deviation.

#### Morphogens, gene regulation and cell fate

Gene expression in the zebrafish hindbrain initially forms a rough r2-6 pattern at 11 hpf, which is refined over time into five precise segments of similar size with four sharp boundaries by 14 hpf (Fig 1C, D). RA synthesized in somites adjacent to the anterior spinal cord diffuses anteriorly and is required for proper rhombomere formation, including direct activation of *vhnf1* in r5 and r6 and *hoxb1a* in r4 [37–39]. Mutual inhibition between *vhnf1* and *irx3* specifies the first pre-rhombomeric r4/r5 boundary at 9.5-10 hpf [48]. RA then activates *hoxb1a* and *vhnf1* which represses *hoxb1a* expression [39], restricting it to r4 where it activates FGF synthesis [41, 45, 55]. FGF diffuses both anteriorly and posteriorly to induce *krox20* in r3 and r5 [42–44]. Through auto-regulation, *krox20* has two steady-state expression levels, either zero or non-zero, depending on the FGF concentration [18, 44], and the r2/r3 and r5/r6 boundaries are specified by *krox20* levels. Auto-regulation and mutual inhibition between *hoxb1a* and *krox20* establish a toggle switch that specifies and refines the r3/r4 and r4/r5 boundaries [17]. As a result, three distinct cell fates are specified by *hoxb1a* and *krox20* expression levels to establish the r2-r6 pattern, specifically, high *hoxb1a* and low *krox20* expression in r4, low *hoxb1a* and high *krox20* expression in r3 and r5, and low expression of both *hoxb1a* and *krox20* in r2 and r6 (Fig 1D).

In the model, morphogens (RA and FGF) were described by stochastic PDEs in a continuum fashion. Regulation of genes downstream of morphogens was modeled using stochastic ODEs for each individual cell. Interactions between morphogens were modeled at a regular rectangular mesh in the morphogen domain, and the downstream genes for each cell were modeled as being located at the center of each individual cell. Numerical interpolation was used to capture the interplay between morphogens and gene regulation modeled at different grid points (**Supplementary Materials S1**).

#### Mechanical models for individual cells

In the cell mechanical model, we used the subcellular element method (SCEM) [16, 56] to model individual cells and cell-cell mechanical interactions involved in cell sorting (**Supplementary Materials S2**). In this computational formalism, an individual cell consists of a constant number of sub-cellular elements (i.e. nodes). Elements interact according to a prescribed force potential. This force between elements within the same cell is repulsive at short ranges and attractive at long ranges to maintain stable cell volume and circular structure. The forces between elements within different cells are repulsive at short ranges to prevent cell overlaps. At longer ranges, the intercellular forces between elements can be either repulsive or attractive depending on cell identities.

#### Cell sorting

In the zebrafish hindbrain, cell sorting has selectivity based on cell identities. One well-known mechanism of selectivity is cell-cell adhesion mediated by Ephrin-Eph signaling. Ephrin-B2 ligands are expressed highly in even-numbered rhombomeres (r2, r4 and r6), while EphA4 receptors are expressed highly in odd-numbered rhombomeres (r3 and r5), and this alternating pattern controls repulsion between cells in one rhombomere and another [52]. Depletion of EphA4 has more dramatic effects on rhombomere boundaries than EphrinB2a, but knockdown of both enhances boundary defects [50]. Krox20 directly activates transcription of *ephA4* [57], thereby regulating Ephrin-Eph mediated cell sorting. Our model mimics the selective cell-cell adhesion between two cells based on their *krox20* expression levels (Fig 1D). Specifically, cells attract each other if they have similar *krox20* levels, and repel each other if one expresses *krox20* and the other does not.

#### Initial conditions

We chose 11 hpf as the starting point for our modeling study soon after the initiation of gene expression in some rhombomeres. At this stage in the simulations cells were assumed to align uniformly in the rectangular tissue domain. For initial gene expression, we first ran simulations with the stochastic gene regulation model over two hours and used those simulation results as the initial condition. We started with equilibrium solution for RA by solving the corresponding steady-state PDE because RA gradients are established as early as 6 hpf [39]. Because *vhnf1* and *irx3* expression appear much earlier than *hoxb1a*, *krox20* and FGF [39], we ran stochastic simulations for RA, *vhnf1* and *irx3* in the first hour with zero values for *vhnf1* and *irx3*. In the second hour, we included stochastic simulations for all morphogens and genes. *krox20* and FGF start with zero expression while *hoxb1a* starts with a constant expression level because *hoxb1a* has weak expression over the hindbrain domain at early stages [17] and shows dynamic changes in expression earlier than FGF or *krox20* [48].

#### One-dimensional gene expression model

In the one-dimensional gene expression models, we consider steady-state solutions of genes and morphogens in a one-dimensional fixed A-P domain (**Supplementary Materials S3**). The initial conditions were chosen similarly to the full model. Equilibrium solutions for RA, *vhnf1* and *irx3* were taken as the initial conditions. Both *krox20* and FGF levels were initially set at zero. We included *hoxb1a* expression at low but non-zero initial levels.

#### A cell sorting-only model

In the model with only cell sorting without gene regulation, cells sorted based on their pre-assigned identities, and cell identities and numbers did not change throughout the simulations. Three cell identities based on threshold levels of high or low expression of *hoxb1a* and *krox20* in the five segments (r2-6) were considered. The initial “salt-and-pepper” pattern of cell identities was sampled by a mixture Gaussian probability distribution based on each cell’s A-P position (**Supplementary Materials S4**).

#### Quantification of cell fate, rhombomere boundary A-P locations and boundary sharpness

Once these three cell identities were determined in r2-r6 we evaluated rhombomere boundaries (**Supplementary Materials S5**). We defined three critical quantities in our simulations: boundary location along the A-P axis, boundary sharpness (represented as a sharpness index, SI) and the number of dislocated cells (DCs) (see **Methods**). Four boundaries between rhombomeres (r2/r3, r3/r4, r4/r5 and r5/r6) are all perpendicular to the A-P axis. For a single boundary determination, we selected a predefined boundary and calculated total deviations of cells located on the wrong side of this predefined boundary. The total deviations were minimized over the A-P position of the predefined boundary. SI was defined by the minima of total deviations and the A-P position of this boundary was defined based on these minima. A lower SI indicates a sharper boundary. DCs are cells located over three cell-diameters away from the rhombomere to which they belong and they are excluded in calculating boundary location. If the number of DCs exceeds 8 cells, we consider the pattern failed.

### Including multiple morphogens and tissue morphogenesis in the models recapitulates the r2-6 pattern

The stochastic multiscale cell-based model successfully recapitulated the dynamics of r2-6 formation observed in the zebrafish hindbrain. As shown for one stochastic simulation with spatial distributions of multiple genes, both RA and FGF signals have noisy distributions over the space (Fig 2C, D). The RA gradient decreases from its origins at the posterior end of the hindbrain to the anterior, while FGF levels are high in r4 where it is secreted and decreases in both anterior and posterior directions. By generating a time series of the spatial patterns of gene expression (Fig 2E-2G), including *hoxb1a*, *krox20, vhnf1 and irx3*, our model recapitulates rhombomere boundary sharpening [17, 49]. For example, *krox20* is weakly expressed in r3 and r5 at 11 hpf and upregulated by 12 hpf, with expression stronger in r3 than r5 (Fig 1A, 2E). At this stage, *hoxb1a*+ and *krox20*+ cells intermingle and a few cells close to the r4/r5 boundary undergo identity switching as they co-express low levels of both *krox20* and *hoxb1a* [49]. By 13 hpf, cells closer to the boundaries and most of the *krox20*/*hoxb1a* co-expressing cells commit to one segment or another and boundaries become sharper. At 14 hpf, all cells segregate to their territories and the boundaries fully sharpen, producing a precise five-segment pattern.

The modeling output naturally accounts for the sharpening of other gene expression boundaries in r2-6, despite differences in their interactions. For example, the anterior edge of *vhnf1* expression and the posterior edge of *irx3* expression specify the position of the pre-rhombomeric r4/r5 boundary at 11 hpf (Fig 2F, G). At later stages, this *vhnf1*/*irx3* border shifts posteriorly to become located in r5 [48]. Unlike *hoxb1a* and *krox20*, *vhnf1* does not auto-induce itself to maintain its own expression without RA signals. Consequently, *vhnf1* shifts posteriorly [39] after 12 hpf as RA decreases everywhere (**Fig S1C**).

Stochastic simulations were repeated independently (Fig 2H). The results are consistent with experimental measurements of rhombomere A-P length at 14 hpf (r3 = 42±5 µm, r4 = 34±5 µm, r5 = 37±4 µm) as well as sharpening. For example, from 11-12 hpf, identity switching affects the sharpness of the r4/r5 boundary [17], with a higher SI and DC number during this period. From 12-14 hpf, SI and DC number gradually decrease to the minimum as all boundaries sharpen.

### Cooperation between RA and FGF improves robustness of initiation of the segmental pattern

A previous model that only considered the RA morphogen gradient without cell sorting or convergent extension successfully simulated many aspects of the formation of the r2-5 pattern [17]. In the model, *hoxb1a* and *krox20* were considered as direct downstream targets of RA, despite *krox20* being indirectly induced by RA through *hoxb1a* and FGF. Our new model incorporates both RA and FGF as well as these additional features of *krox20* regulation.

Comparing the two-morphogen (RA + FGF) to the one-morphogen (RA) model reveals many similarities and some key differences (Fig 3). In both, the borders between *hoxb1a* and *krox20* specify r3/r4 and r4/r5 boundaries, the border between *vhnf1* and *irx3* lies posterior to the r4/r5 boundary, and *krox20* has two steady-state levels induced by either RA (one-morphogen model) or FGF (two-morphogen model) depending on the morphogen levels. In the two-morphogen model, FGF has the highest expression in r4 where it is secreted and decreases in both anterior and posterior directions. By inducing *krox20* in a concentration-dependent manner, FGF can specify the r2/r3 and r5/r6 boundaries. In contrast, the one-morphogen model, with RA decreasing monotonically from posterior to anterior, can only specify r2/r3 and not the r5/r6 boundary. Overall, the two-morphogen model can specify four boundaries (r2/r3, r3/r4, r4/r5 and r5/r6), while the one-morphogen model can specify only three boundaries (r2/r3, r3/r4 and r4/r5).

**Fig 3.**
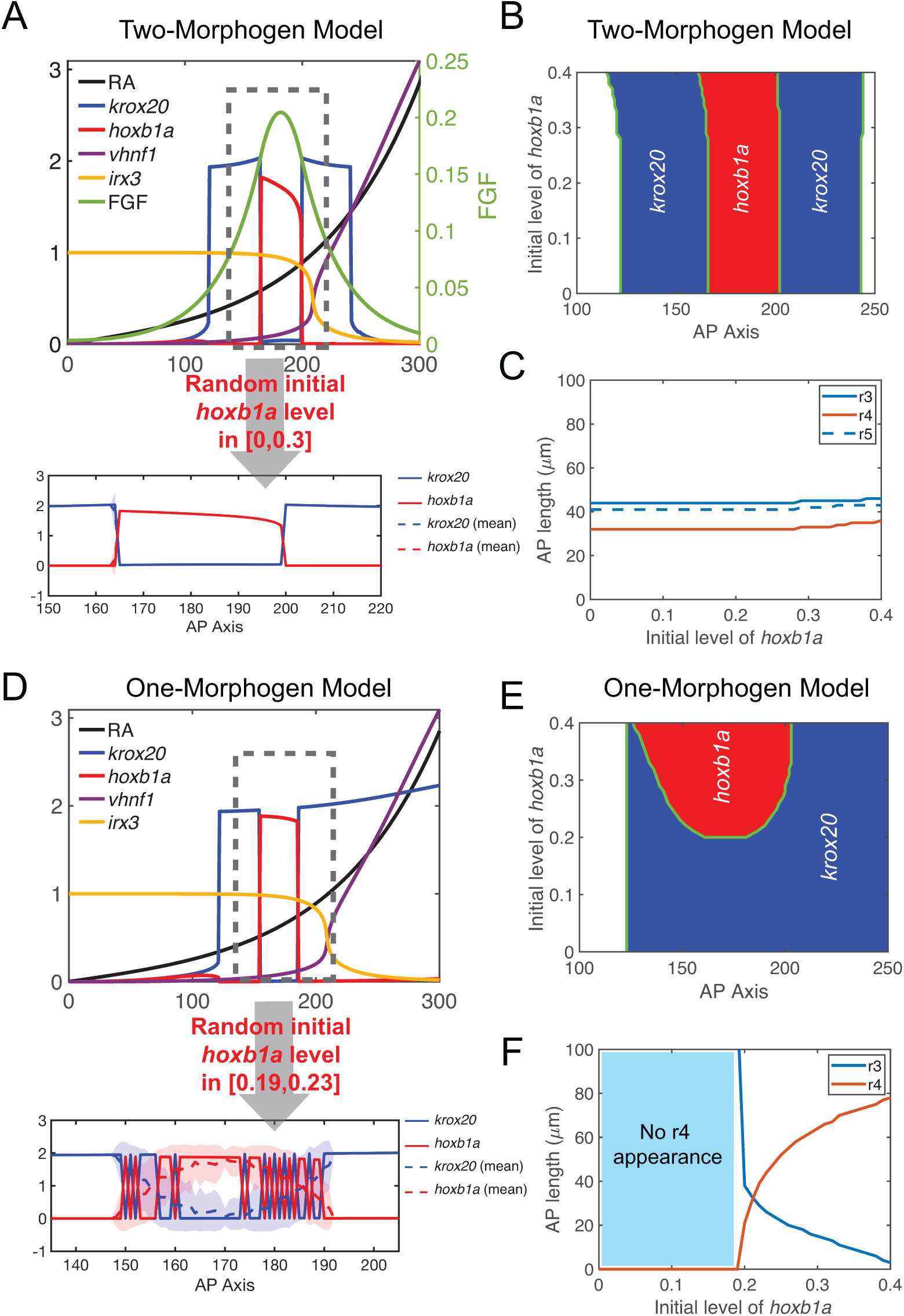
Comparing two-morphogen (RA, FGF) and one-morphogen (RA) models. **(A-C)** One-dimensional simulations for the two-morphogen model. **(A)** The upper panel shows spatial distributions of RA, *krox20*, *hoxb1a*, *vhnf1*, *irx3* and FGF. The initial *hoxb1a* level is modeled as a constant 0.21 over the space. In the lower panel, the initial *hoxb1a* level is randomly sampled over the space independent of the location. The value is randomly uniformly distributed at a level of [0,0.3]. Solid line represents one simulation. Dashed line represents average values and the width of the shading around each line represents the standard deviation (n=100). Since fluctuations over multiple simulations are small, solid lines overlap with dashed lines and the small standard deviations result in shadings of negligible width around the dashed lines. X-axis, microns; Y-axis, arbitrary units. **(B)** Phase diagram of *hoxb1a* and *krox20* distributions with different initial *hoxb1a* levels. **(C)** Rhombomere lengths with different initial *hoxb1a* levels. **(D-F)** Similar one-dimensional simulations for the one-morphogen model. For **(D)**, in the upper panel, the constant initial *hoxb1a* level is taken as 0.21; in the lower panel, the initial *hoxb1a* level is randomly sampled with levels in the range [0.19,0.23] with uniform distribution. Corresponding (**E**) phase diagram and (**F**) graph of rhombomere lengths with the one-morphogen model.

Additionally, we compared robustness in the two models with respect to initial *hoxb1a* expression. With constant initial *hoxb1a* level everywhere in space, we compared phase diagrams and resulting rhombomere lengths between the two-morphogen and one-morphogen models (Fig 3B, C, E, F). Interestingly, the inclusion of FGF makes the model relatively less sensitive to initial *hoxb1a* levels, in terms of the locations of gene expression boundaries and sizes of r3-5. When the initial *hoxb1a* level exceeds a certain level (e.g. 0.3), three rhombomeres (r3, r4 and r5) expand slightly along the A-P axis with r2/r3 and r3/r4 expanding anteriorly, and r4/r5 and r5/r6 expanding posteriorly (Fig 3B, C). In contrast, with RA alone, lengths of r3 and r4 are more sensitive to initial *hoxb1a* levels. In simulations with low initial *hoxb1a* levels (<0.2), r4 does not form (Fig 3E, F). As initial *hoxb1a* levels increase from 0.2 to 0.4, the r4 region rapidly expands at the expense of r3. A 15% increase in initial *hoxb1a* level (0.2-0.23) leads to an over two-fold expansion of r4 (21-44 µm) and r3 essentially vanishes when the initial *hoxb1a* level is close to 0.4. Thus, the two-morphogen model outperforms the one-morphogen model in robustness of rhombomere length, in that the second morphogen buffers responses to initial gene expression variation.

We also examined how the pattern reacts to perturbations of initial gene expression. We consider noisy initial *hoxb1a* levels over the space. In the two-morphogen model, such noise has negligible effects on later *hoxb1a* and *krox20* distributions resulting in clear segmental patterns and sharp r3/r4 and r4/r5 boundaries, and multiple simulations result in almost identical patterns (Fig 3A). However, in the one-morphogen model, *hoxb1a* and *krox20* distributions fluctuate more dramatically than the two-morphogen model. Despite a much smaller magnitude of perturbation (13%) in initial *hoxb1a* levels, the one-morphogen model shows fluctuating boundaries for both r3/r4 and r4/r5 (Fig 3D). Overall, the two-morphogen cooperation facilitates both accurate rhombomere lengths and sharp boundaries with perturbations of initial gene expression, providing robustness in patterning during the initial stages.

### Rapid initial convergence improves boundary sharpness and segment size

Due to stochasticity, the initial pattern shows rough boundaries between rhombomeres. Later on, patterns sharpen and refine boundaries from 11-14 hpf. During these stages, the zebrafish hindbrain changes shape dramatically as it narrows in width along the L-R axis, extends in length along the A-P axis (Fig 1A) and thickens in the D-V axis [58]. We mainly studied A-P and L-R axes and their dimensions were experimentally measured (Fig 2A, 2B). Length along the A-P axis changes slowly during this period, but width along the L-R axis rapidly shrinks from 283 µm to 162 µm during the first hour, and further to 104 µm during the last two hours with approximately a 75% drop in the average rate after which patterning is largely complete. To determine how such rapid initial convergence influences hindbrain patterning we compared models incorporating medium or slow initial convergence with the rapid initial convergence rate we measured experimentally. All three types of convergence have the same initial and terminal L-R width. The curve of the medium convergence is taken as a linear function. The curves of the slow and rapid initial convergences are symmetric to the linear curve (Fig 4A).

**Fig 4.**
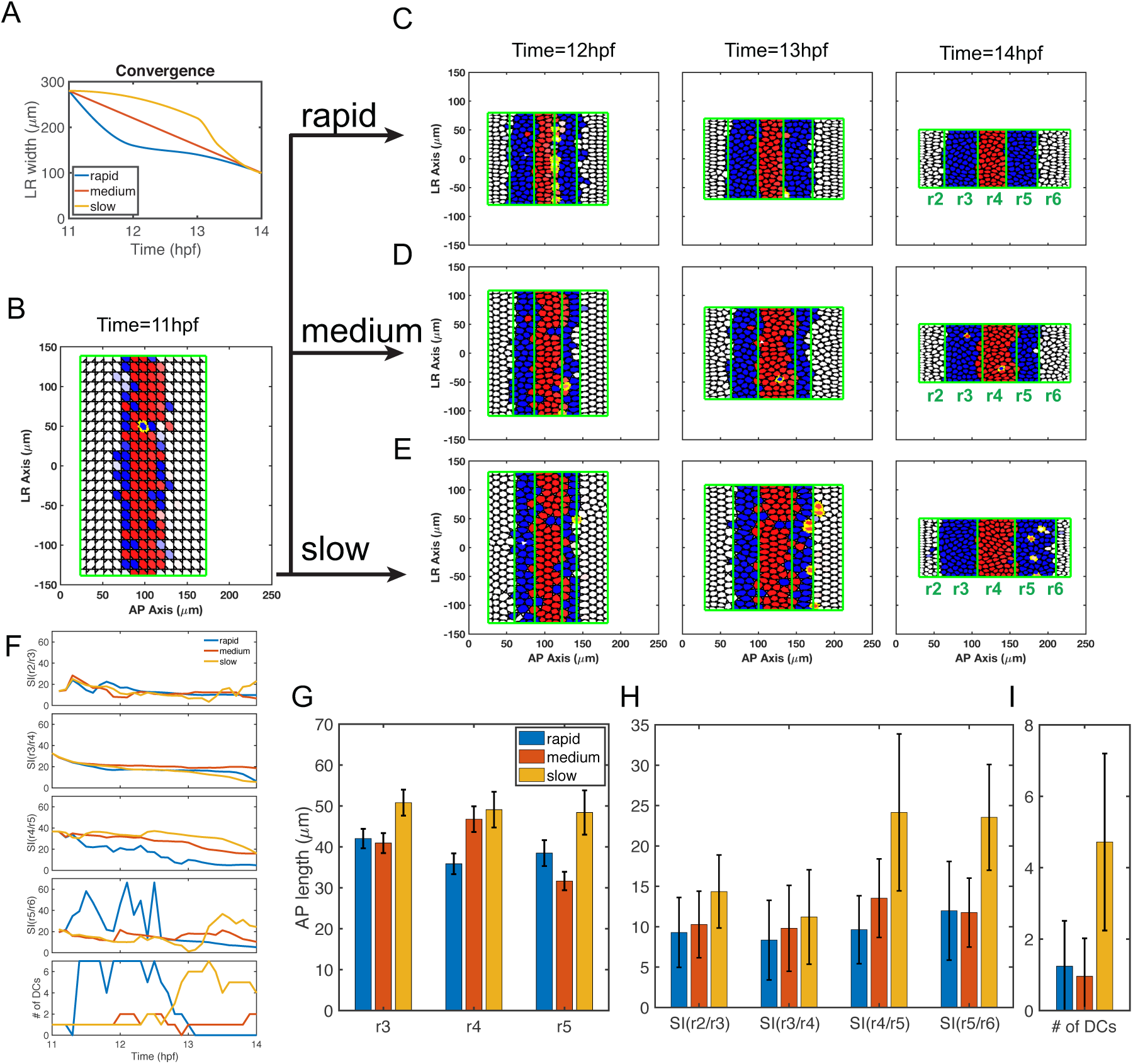
Simulations of full models combining gene regulation and cell sorting with different convergence rates. **(A)** Three convergence rates during the 11-14 hpf period are considered in the model, rapid (from experimental measurements, Fig 2B), medium and slow. All start and terminate with the same L-R width. The curve of medium convergence is depicted as a linear function. The curve of slow initial convergence is symmetric to the curve of rapid initial convergence with respect to the curve of linear function. **(B-E)** Time series of cell distributions with different convergence rates from 11-14 hpf. *hoxb1a* (red) and *krox20* (blue) expression. Dislocated cells (DCs) are highlighted by yellow edges. **(B)** Three simulations start with the same initial cell distribution (11 hpf) generated by the gene expression model (see Methods). Cell distributions with **(C)** rapid, **(D)** medium and **(E)** slow initial convergence rates from 12-14 hpf. **(F)** The boundary sharpness index (SI) for four boundaries (SI(r2/r3), SI(r3/r4), SI(r4/r5) and SI(r5/r6)) and DC number versus time. **(G-I)** Histograms depicting three convergence rates analyzed for **(G)** rhombomere lengths of r3-5, **(H)** SI and **(I)** DC number. Each represents 100 independent stochastic simulations for each convergence rate based on the same parameters. Error bars represent standard deviation.

We first performed simulations with full models including both gene regulation and cell sorting. Regardless of the convergence speed, most cells segregate to their correct territories and the final patterns display sharp boundaries between rhombomeres (Fig 4B-4E). Rapid initial convergence allows the sharpest boundaries and the fewest DCs while slow initial convergence results in the roughest boundaries and more DCs (Fig 4F). SI and DC numbers at 14 hpf in multiple independent simulations confirm conclusions based on single simulations (Fig 4H, I). The exception is that rapid and medium convergence rates yield similar SI and DC numbers at the r5/r6 boundary (Fig 4H, I). Rhombomere A-P lengths also vary greatly in this model with different convergence rates (Fig 4G). Simulations with slow initial convergence result in all three rhombomeres (r3-5) elongated. Simulations with medium convergence result in a shorter r5 compared to r3 and r4. Simulations with rapid initial convergence result in r3-5 being all roughly the same length (with r4 slightly shorter) similar to experimental measurements of the hindbrain at 14 hpf (Fig 2H, 4G).

Taken together, rapid initial convergence facilitates robust patterns by optimizing boundary sharpness and rhombomere A-P length. These influences depend on multiple mechanisms, including cell sorting and gene regulation, and their coordination.

### Rapid initial convergence improves boundary sharpness through cell sorting

Next, we performed simulations with models incorporating cell sorting alone and excluding gene regulation using the cell sorting-only model. Initial “salt-and-pepper” cell distributions by assigning each cell an identity generated the five-segment pattern with rough boundaries (Fig 5A). Similar to observations in full models (Fig 4), most cells segregate to their correct territories and the final patterns display sharp boundaries between rhombomeres (Fig 5B-D). Rapid initial convergence allows the sharpest boundaries and the fewest DCs, while slow initial convergence results in the roughest boundaries and more DCs (Fig 5E). SI and DC numbers at 14 hpf in multiple independent simulations confirm conclusions based on single simulations (Fig 5G, H). In the model with cell sorting alone, while these different speeds of convergence have major effects on boundary sharpness, they have relatively minor effects on rhombomere A-P length (Fig 5F). These results suggest rapid initial convergence improves boundary sharpness by facilitating the efficiency of cell sorting.

**Fig 5.**
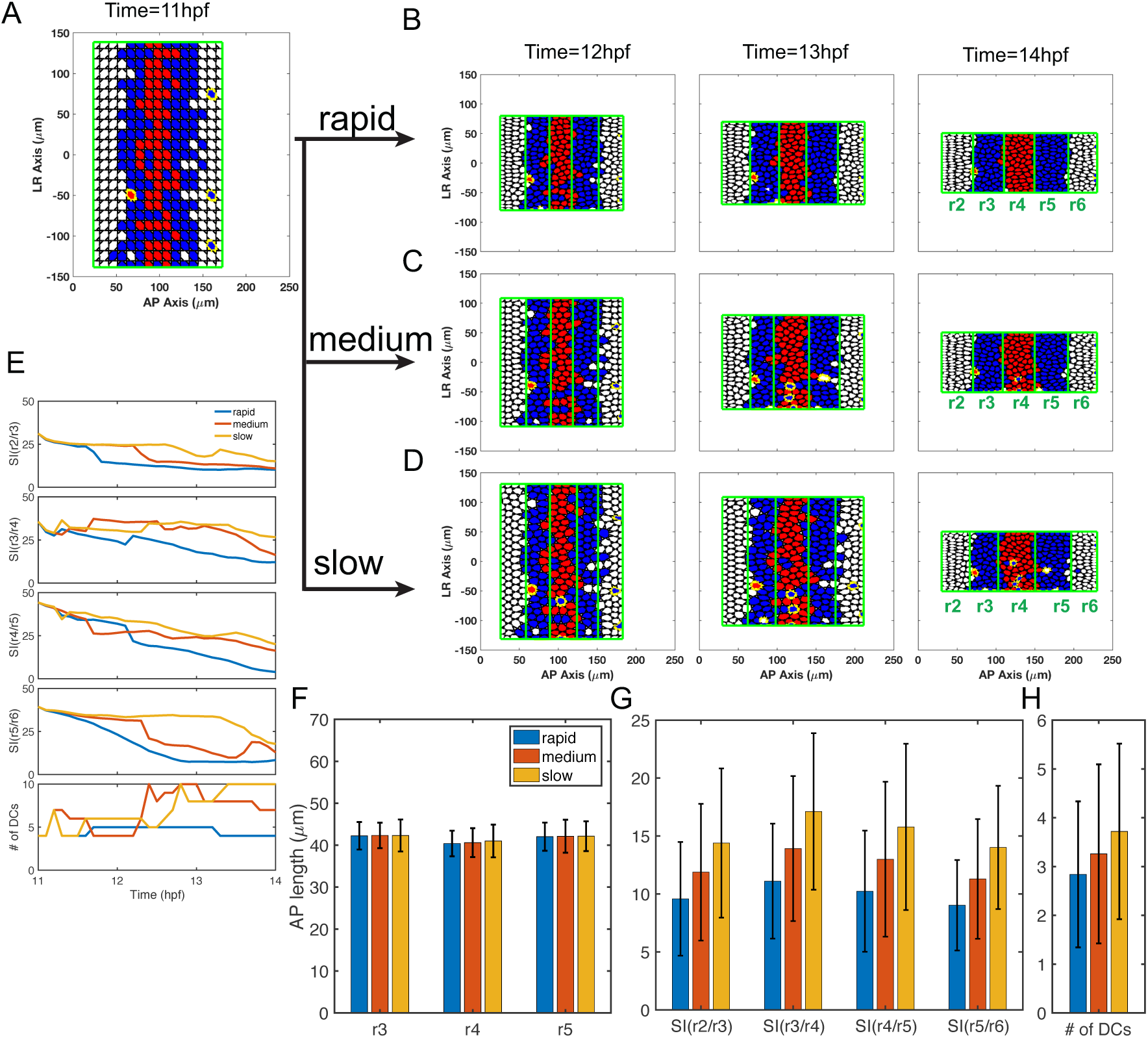
Simulations with selective cell-cell adhesion/sorting alone with different convergence rates. **(A-D)** Time series of cell distributions with different convergence rates from 11 to 14 hpf. *hoxb1a* (red) and *krox20* (blue) expression. Dislocated cells (DCs) are highlighted by yellow edges. **(A)** Three simulations start with the same initial cell distribution (11 hpf) generated by the Gaussian mixture distribution. Cell distributions with **(B)** rapid, **(C)** medium and **(D)** slow initial convergence from 12-14 hpf. **(E)** The boundary sharpness index (SI) for four boundaries (SI(r2/r3), SI(r3/r4), SI(r4/r5) and SI(r5/r6)) and number of DCs versus time. **(F-H)** Histograms depicted three convergence rates analyzed for their **(F)** rhombomere lengths of r3-5, **(G)** SI and **(H)** the DC number. Each represents 100 independent stochastic simulations for each convergence rate are based on the same parameters. Error bars represent standard deviation.

### Rapid initial convergence helps specify correct rhombomere lengths by regulating cell fate

To investigate the effects of convergence on rhombomere lengths, we studied dynamics of gene expression under different convergence rates. We first investigated the influence of convergence rate on spatiotemporal dynamics of morphogen distribution and cell fate commitment (Fig 6). With rapid initial convergence, the RA signal and its direct target, *vhnf1*, increase quickly from 11-12 hpf, particularly in the posterior hindbrain (150 µm and 200 µm in Fig 6A, B), then decrease gradually. Near the r4/r5 boundary, *hoxb1a* is repressed by the increasing levels of *vhnf1*. FGF made in r4 then activates *krox20* leading to identity switching for cells near the r4/r5 boundary with low *hoxb1a* expression (Fig 4C). With medium or slow initial convergence, the RA signal and *vhnf1* remain relatively unchanged compared to rapid initial convergence (Fig 6A, B). Indeed, we observed that fewer cells switch from an r4 to an r5 identity and the r4/r5 boundary is located further posteriorly with medium or slow initial convergence (Fig 4C-E). Similar to RA, FGF levels increase and peak around 12 hpf with models that include rapid initial convergence (Fig 6C). With medium convergence, FGF levels remain relatively unchanged, while with slow initial convergence, FGF levels remain unchanged at the early stage then increase and peak quickly at 13-14 hpf. At the same A-P position, slow initial convergence results in higher maximum FGF levels than rapid or medium convergence rates. Since FGF induces *krox20* expression to drive identity switching from r2 identity to r3 (and r6 to r5) identity, the higher maximum FGF levels that result from slow initial convergence lead to displacement of the r2/r3 boundary anteriorly and the r5/6 boundary further posteriorly than rapid or medium convergence rates.

**Fig 6.**
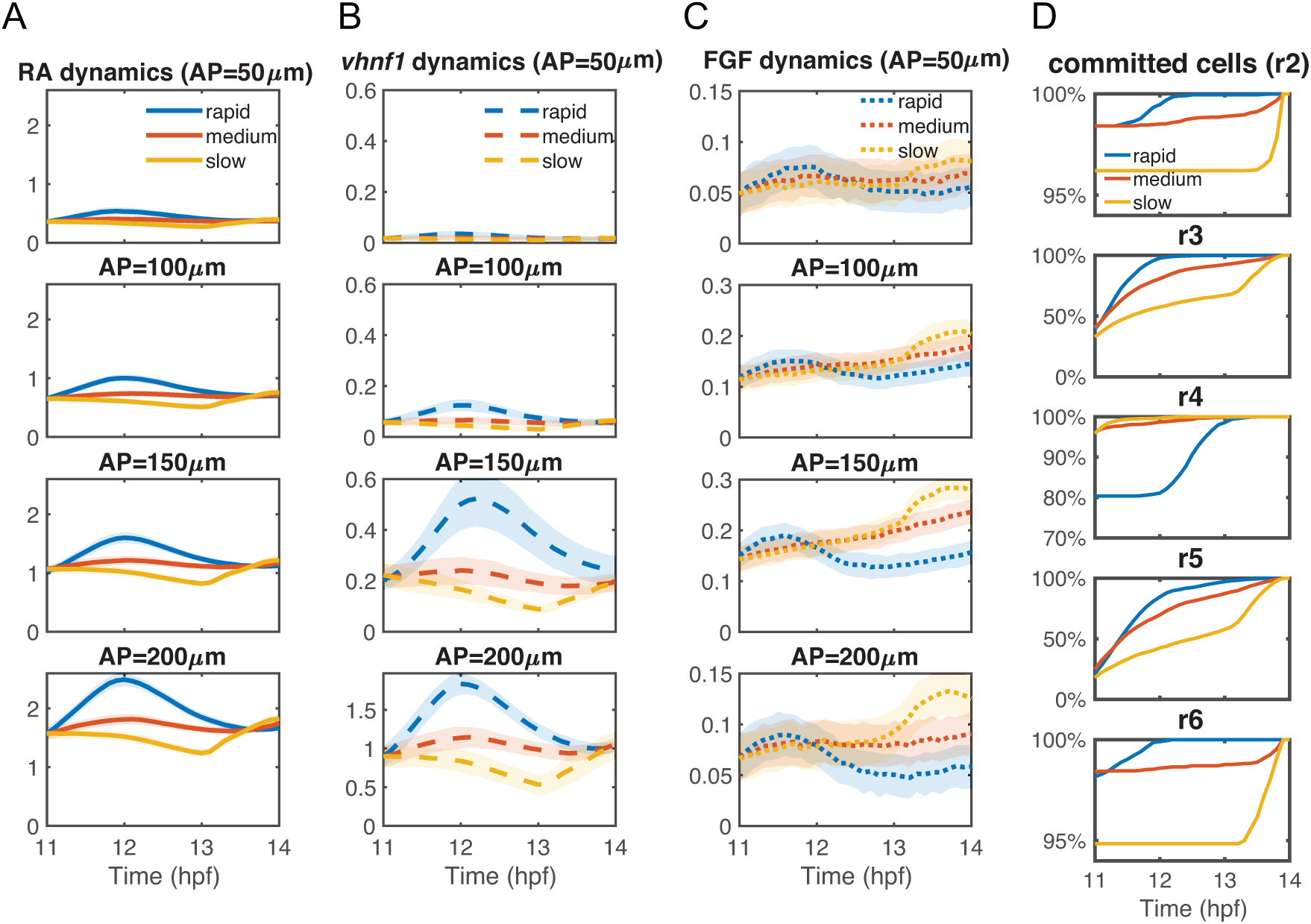
Dynamics of morphogens and cell commitment time with different convergence rates. The statistics of the dynamics of **(A)** intracellular RA [*RA*]*_in_*, **(B)** *vhnf1*, and **(C)** FGF signaling *[FGF]_signal_* at different A-P lengths of the tissue domain: 50 µm, 100 µm, 150 µm and 200 µm. Lines represent average values and the width of the shading around each line represents the standard deviation. **(D)** The temporal dynamics of total percentages of cells that have committed in each rhombomere (r2-r6). Each panel shows the dynamics of the percentages of cells that have committed in each rhombomere. Data are collected from the full models, see Fig 4.

The different rhombomere A-P lengths under different convergence rates can be observed consistently either in the full model (Fig 4) or in the model excluding cell sorting (**Fig S2**). The dynamics of the morphogens provides an explanation for the length behaviors. Rapid initial convergence leads to the smallest r4 because more r4 cells switch to r5 identities near the r4/r5 boundary. Slow initial convergence leads to a larger r3 and r5 because more r2 or r6 cells switch to r3 or r5 identities. Medium convergence results in the smallest r5 because fewer cells switch near the r4/r5 boundary than rapid initial convergence and fewer cells switch near the r5/r6 boundary than with slow initial convergence.

Interestingly, r3 emerges earlier and it initially has a larger A-P length than r5 since *vhnf1* expressed posteriorly represses the FGF activator, *hoxb1a*, resulting in weaker FGF signaling in r5 (Fig 4A). Through rapid convergence, r4 cells can switch to an r5 identity to compensate for the difference between r3 and r5 lengths to achieve correct A-P rhombomere lengths similar to experimental measurements.

### Rapid initial convergence improves boundary sharpness through synergy between gene regulation and cell sorting

To study potential coordination between gene regulation and cell sorting in boundary formation, we restricted them individually at different patterning stages. Minimizing gene regulatory interactions in the model at early stages causes major patterning defects, while at later stages effects on boundary sharpness are minor (**Fig S3**). Conversely, limiting selective cell sorting to one-hour intervals within 11-14 hpf (with cells allowed to sort uniformly at other times) yields the worst patterns when limited to 11-12 hpf (**Fig S4**). These results suggest that gene regulation is more important during early patterning stages and cell sorting later for boundary sharpening.

As shown in the previous subsection, with rapid initial convergence rates morphogen levels increase and peak at around 12 hpf, while morphogen levels increase and peak much later with medium or slow initial convergence. Since *hoxb1a* and *krox20* are direct targets of RA and FGF, respectively, the timing of cell commitment as measured by expression of these genes is closely tied to the timing of peak morphogen levels. Consequently, rapid initial convergence drives the earliest cell commitment at around 12 hpf. Slow initial convergence leads to much later cell commitment (Fig 6D). One notable exception is that r4 cells commit later in the case of rapid initial convergence, due to switching near the r4/r5 boundary, but most cells still commit before 13 hpf (Fig 6D). Driven by early cell commitment, rapid initial convergence extends the effective period of cell sorting and shortens the effective period of gene regulation to improve boundary sharpness.

### Rapid initial convergence mediates the trade-off between rhombomere length and boundary sharpness

To examine the sensitivity of our observations to model parameters, we performed a large number of simulations with random parameters, assaying both rhombomere A-P length and boundary sharpness. Using n=1000 independent repeats for each convergence rate, we found 513, 563 and 452 simulations that successfully produced a five-segment pattern for rapid, medium and slow initial convergence, respectively.

These results also revealed a trade-off between rhombomere length and sharpness, i.e. reduced length typically resulted in higher Sis, indicating rougher boundaries (Fig 7 and **Fig S5**). This can likely be explained by the fact that a shorter rhombomere has fewer cells with the same identity, thus cell sorting has less effect and cells are more susceptible to noise. As a result, a single stray cell has more impact on the boundary sharpness index. Such trade-offs are also observed for the model without convergent extension, leading to shorter and rougher rhombomere lengths (**Supplementary Material S11)**. With shorter rhombomeres, rapid initial convergence significantly reduces this trade-off, particularly in r4 and r5 (Fig 7 and **Fig S5**).

**Fig 7.**
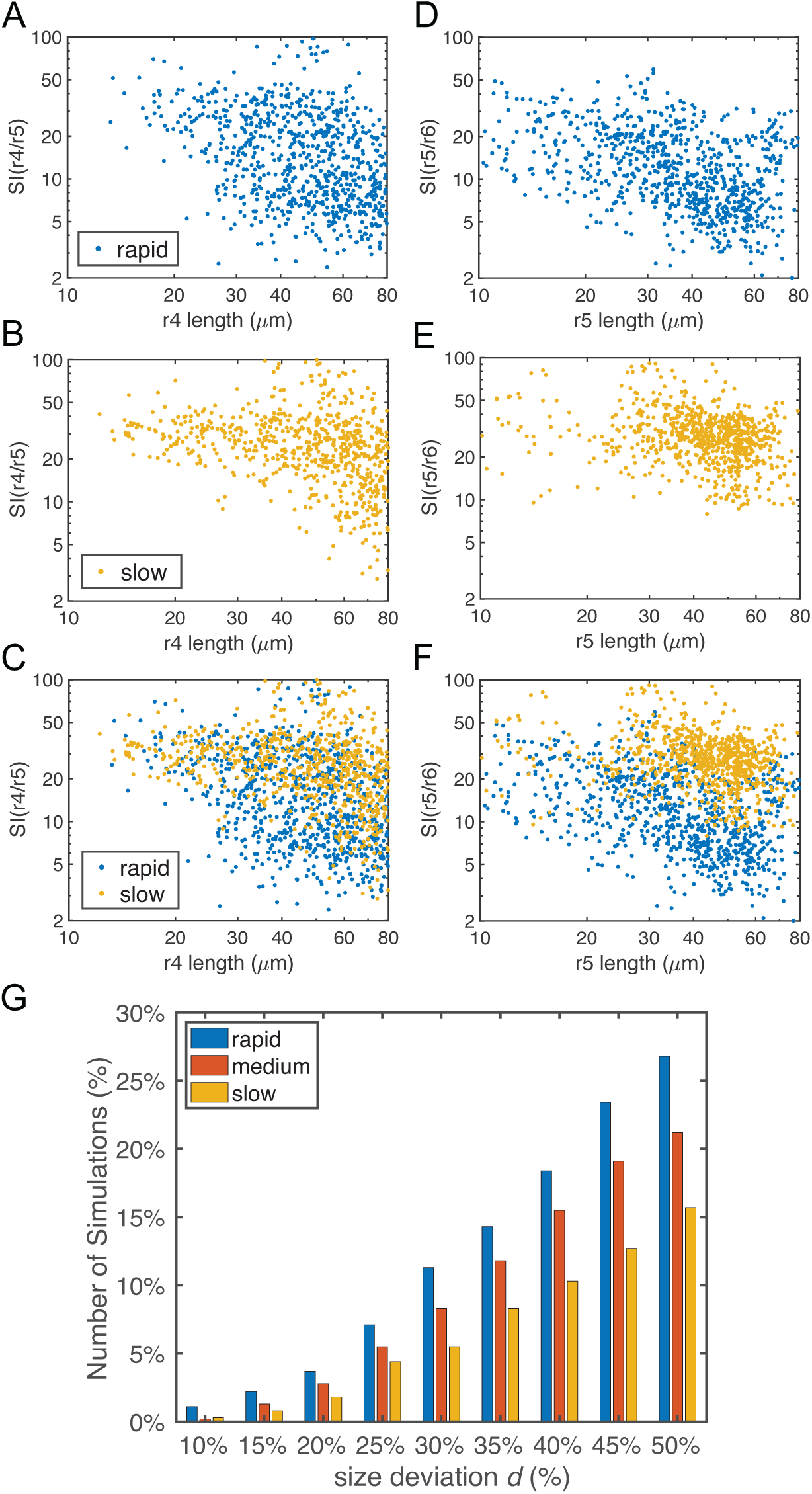
Boundary sharpness and rhombomere lengths based on simulations with random parameters in gene regulation. Parameters for gene regulation were randomly perturbed and a total of n=1000 simulations are displayed for each convergence rate. There are 513, 563 and 452 simulations for rapid, medium and slow initial convergence, respectively, which successfully generate the r2-6 pattern with four boundaries. **(A-F)** Dot plots showing the relationship between rhombomere length and boundary sharpness. Each point represents the corresponding quantities for each simulation. **(A-C)** Length of r4 versus sharpness index (SI) of the r4/r5 boundary with: **(A)** rapid initial convergence, **(B)** slow initial convergence and **(C)** a comparison between rapid and slow initial convergence. **(D-F)** Length of r5 versus SI of r5/r6 boundary with: **(D)** rapid initial convergence, **(E)** slow initial convergence and **(F)** a comparison between rapid and slow initial convergence. **(G)** Fractions of simulations achieving roughly equal rhombomere lengths versus the deviation *d%*. With a deviation *d*, a simulation has roughly equal rhombomere lengths if the length of each rhombomere is deviated at most *d*% from its average experimental length (i.e. r3, r4 and r5 are in the ranges of 42*(100%±*d*%) µm, 34*(100%±*d*%) µm and 37*(100%±*d*%) µm, respectively).

Moreover, we quantified fractions of simulations achieving roughly equal segment length. We considered a simulation having equal rhombomere lengths if A-P lengths of r3, r4 and r5 were close to their experimentally measured average lengths within ranges[*m* * (100% −*d*%), *m* * (100% + *d*%)], where *m* was the measured average length (**Table S7**). With any values of deviation *d*, there are higher fractions of simulations achieving roughly equal length under rapid initial convergence compared to medium and slow initial convergence at 14 hpf (Fig 7G). Experimentally, standard deviations of r3, r4 and r5 length are within the range between 10% and 15% (**Table S7**). Within this range of *d*, rapid initial convergence has at least 69% and 175% higher fractions of simulations achieving roughly equal segment length than medium and slow initial convergence, respectively (Fig 7G). These results are consistent with our findings that rapid initial convergence generates more accurate lengths of rhombomere and sharper boundaries.

## DISCUSSION

Our models suggest that a combination of two morphogens and rapid initial tissue convergence together drive robust hindbrain segmentation. Inclusion in the model of the short-range morphogen (FGF) secreted from r4, combined with the longer-range morphogen (RA) secreted posteriorly, substantially improves the robustness of segmental patterning compared with RA alone. Cooperation between morphogens is common in pattern formation in many contexts, in part because it helps maintain accuracy in size and boundary sharpness of target gene expression domains. Our previous models and experiments in the hindbrain have focused primarily on the r4/r5 boundary, where many gene regulatory interactions are known and the RA gradient is relatively steep [17, 49, 50]. The current model expands upon this work to explain the formation of other rhombomere boundaries, particularly r2/r3, r3/r4 and r5/r6, with the additional positional information provided by FGF. Surprisingly, rapid initial convergence dramatically improves robustness of rhombomere patterning, both segment size and boundary sharpness. Rapid initial convergence may also be a conserved strategy for precise establishment of gene expression domains in other embryonic tissues that elongate by convergent extension [59–61] such as axial mesoderm in early vertebrate embryos [62] or stacking of chondrocytes in developing cartilages [63, 64].

### Complementary roles of long- and short-range morphogens in pattern accuracy and precision

In morphogen gradient-mediated patterning, it is crucial not only that target gene expression boundaries are accurately positioned but also that they are sharp. However, there is a trade-off between accuracy and precision of boundary patterning that depends on morphogen gradient steepness [1, 27]. A steep morphogen gradient specifies boundary locations more precisely in the face of fluctuations in signal, is less sensitive to noise and facilitates boundary sharpness. However, the trade-off is that it makes positioning boundaries more sensitive to perturbations or noise in morphogen synthesis, slight shifts in which can move the boundaries along the A-P axis. Since RA is responsible for the initial A-P patterning of the hindbrain, starting from gastrulation, it likely plays a more prominent role in accuracy, due to its shallow distribution across much of the patterning region [40] and self-enhanced degradation [28]. On the other hand, FGF likely plays a prominent role in precision to help improve the sharpness of boundaries adjacent to its source, since its gradients are likely steeper due to its local effects [41, 45, 55]. In addition, FGF synthesis most likely varies less since one of its upstream regulators, *hoxb1a*, is bi-stable and tightly controlled by a complex network [46, 47, 65]. Together, these complementary features of the long-range shallow RA gradient and the short-range steep FGF gradient help overcome the trade-offs inherent in morphogen patterning systems for achieving both accurate and precise rhombomere pattern.

During hindbrain segmentation, r4 becomes the secondary signaling center that produces FGFs (e.g. Fgf4 and Fgf8) in zebrafish [41, 42, 45, 55] that are regulated by the posteriorizing signal RA [28]. The MHB is another secondary FGF (i.e. Fgf8 in zebrafish) signaling center that likely contributes to anterior rhombomere patterning [66]. In many biological contexts, two morphogens interact with each other to facilitate spatial pattern formation. Interactions between the long- and short-range morphogens induce Turing patterns, such as Sox9 and Bmp in digit patterning [12], Edar and FGF in murine tooth development [67], FGF and Shh in limb regeneration [68], and Nodal and Lefty in early mesoderm formation and left-right patterning [69, 70]. Two long-range morphogens with anti-parallel distributions improve the precision of a single boundary, such as Bcd and Cad in *Drosophila* embryo segmentation [71], and Bmp and Shh in vertebrate neural tube patterning [14, 71, 72]. Unlike these examples, the novel two-morphogen mechanism presented in this work includes one long-range and one short-range morphogen that act in parallel on downstream targets. This system specifies multiple boundaries of gene expression and improves both accuracy and precision of segmental patterns.

### Rapid initial convergence in tissue morphogenesis improves pattern robustness

Intuitively, elongation along the A-P axis might be expected to hinder segmental patterning and rhombomere boundary sharpening [17], since cells quickly change neighbors and intercalate. However, we find quite the opposite (Fig 8). Rapid initial convergence facilitates boundary sharpening through two strategies. First, it induces stronger intercellular interactions, consequently stronger cell sorting, leading to sharper boundaries (Fig 8A). Second, it induces an early peak of morphogens that can result in early cell commitment, allowing cell sorting sufficient time for rearrangements without disrupting cell fate switching (Fig 8B). Rapid initial convergence can also regulate rhombomere length through morphogen dynamics. Initially the length of r5 is shorter than r3. Through a steeper RA distribution induced by rapid initial convergence, cells switch from an r4 to an r5 leading to similar r3 and r5 lengths (Fig 8C).

**Fig 8.**
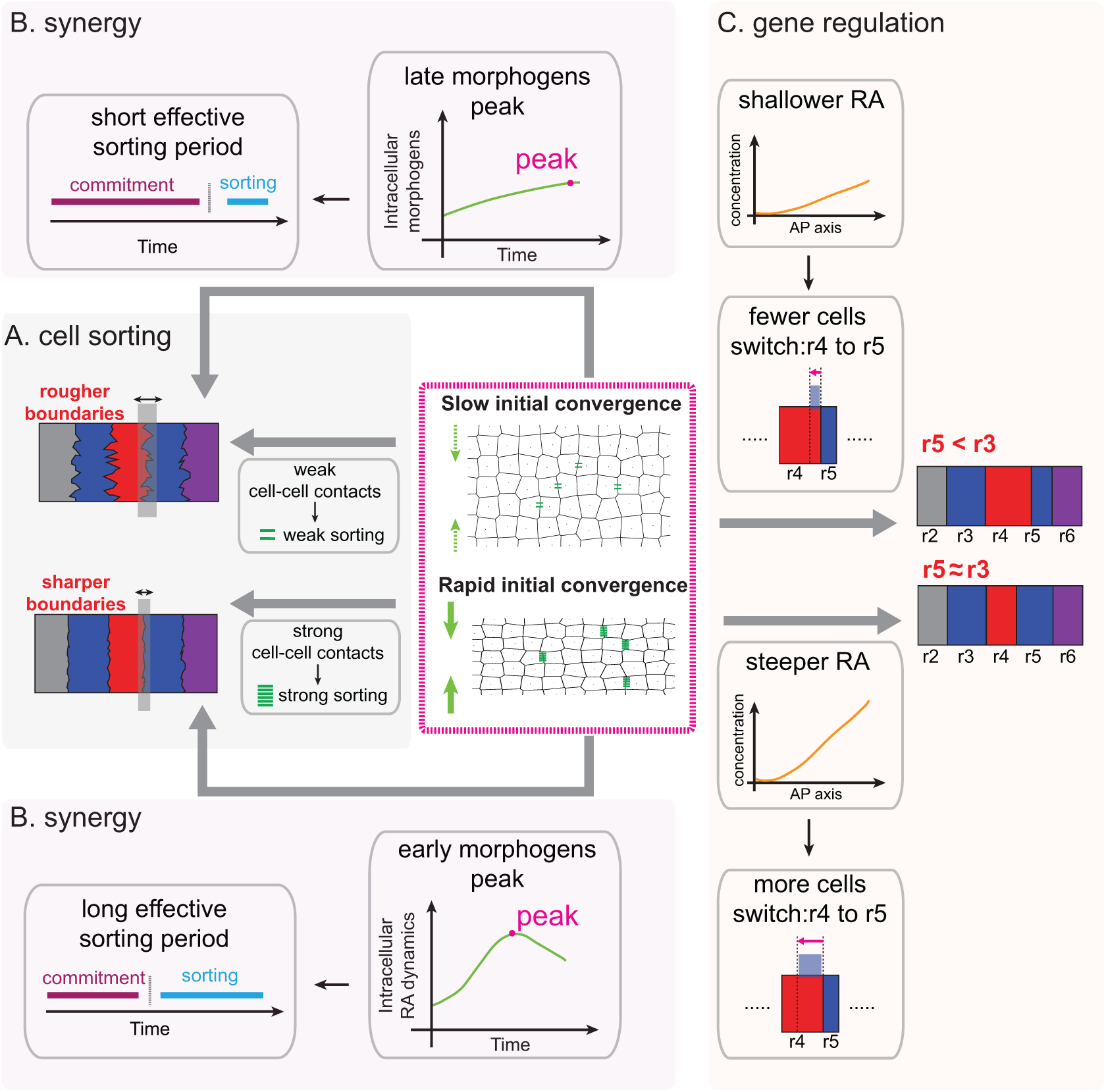
Schematic illustrating how rapid initial convergence improves pattern robustness. **(A)** Cell sorting: rapid initial convergence increases cell-cell contacts to enhance strength of cell sorting, leading to sharper boundaries. Number of green lines represent strength of cell sorting. **(B)** Synergy between cell sorting and gene regulation: rapid initial convergence induces an early peak of morphogens for both RA and FGF, leading to an early commitment of cell fates. Cell sorting mechanisms fully function to sharpen boundaries with sufficient time without disrupting cell fate switching. **(C)** Gene regulation/cell fate: rapid initial convergence produces a steeper RA distribution to induce more cells switching from r4 (red) to r5 (blue) identity. Consequently, r5 has similar size with r3, consistent with the experimental measurements.

Another consequence of convergent extension is the movement of morphogen production and responding cells relative to one another. RA levels increase with time during early stages of hindbrain development due to increased synthesis and accumulation [39], then decrease due to the movement of the source of RA (*aldha1a* expression) further posteriorly as the body axis elongates [39, 46, 47]. Our model successfully recapitulates these RA dynamics (Fig 6A), which are also critical for specifying rhombomeres of the correct length and boundary sharpness. Previous manipulations of convergent extension in early zebrafish or mouse embryos, which result in a shortened body axis, have shown that convergent extension is critical in establishing signaling gradients and subsequently maintaining robust segmental patterning of the hindbrain, consistent with our results [73, 74].

As tissue deforms, extracellular morphogens may have both active motion driven by the tissue dynamics as well as movements of the signals induced by morphogens within cells. In our model, both extra and intracellular morphogens were modeled in continuum context with Eulerian coordinates, where the advections are usually required to capture morphogen dynamics with moving boundaries [75]. We found that even if we removed intracellular advections from the models, our main results showing a positive influence of rapid initial convergence on patterning remain (**Fig S11**).

### Tissue size, thickening and additional signals in hindbrain patterning

The embryonic zebrafish hindbrain is extremely small and composed of relatively few cells compared with most other vertebrates [49, 76, 77]. The actual A-P length of each rhombomere at the stages we have examined (11-14 hpf) is approximately 3∼5 cell-diameters. This small size presents a challenge for sharpening rhombomere boundaries, where a few neighbor cells with the same identity provide weak adhesion during sorting, and even more so for generating a series of rhombomeres of similar size. A rapid initial convergence rate may be particularly important for coordinating size and boundary sharpness in such a miniaturized embryo. However, given the conserved patterns of gene expression and neuronal differentiation observed in hindbrains across species, we are confident that many of the same rules apply.

Our modeling and experimental measurements correspond in many respects, including the dynamics of RA synthesis and FGF4/8 expression as well as *hoxb1a*, *krox20*, *vhnf1* and *irx3* expression in zebrafish. However, many questions remain. In our simulations, DCs remain in r2 and r6 due to randomness in gene expression. For example, in some cases *hoxb1a+* cells are observed in r2 because of early increases in RA, which induce *hoxb1a*. The model cannot account for how these cells switch to their correct segmental identities, but perhaps they are displaced along the D-V axis, undergo apoptosis or are extruded from the hindbrain. Such switching may also reflect a “community effect” by which cells switch identity depending on the collective influences of their neighbors, but the underlying mechanisms have not been fully identified [34, 49]. Gene regulatory networks with other signals involved may also prevent cell switching or sorting by regulating *hoxb1a* or *krox20*. For example, Fgf8 is expressed at the MHB in zebrafish embryos starting at 10 hpf, and likely important for the patterning of more anterior rhombomeres, r1-3, which will be interesting to consider in future models [47, 66]. Wnt is another morphogen that controls early anterior rhombomeres and MHB formation [47, 78].

Many other features of tissue morphogenesis also need to be considered for a comprehensive three-dimensional model of hindbrain segmentation. During the patterning period considered here, cells divide and the NP thickens along the D-V axis [58]. While cells divide in this period, the NP thickens and cell number in the two-dimensional plane (A-P and L-R plane) changes very little [58]. We also studied a two-dimensional model that incorporates cell proliferation and growth and while this led to tighter cell distributions and higher variations in rhombomere length than in other models, overall it confirmed that rapid initial convergence improves pattern robustness (**Supplementary Material S10)**. The two-dimensional nature of our models, which do not consider the complicated dynamics associated with proliferation and thickening along the D-V axis, likely explains why the computed rhombomere lengths in our model do not perfectly fit the experimental measurements (Fig 2H). More realistic, three-dimensional models that incorporate these components pose an exciting challenge and opportunity for the future.

## Author contributions

**Conceptualization**: YQ, LF, TFS and QN

**Data Curation**: YQ, LF, TFS and QN

**Formal Analysis**: YQ, LF, TFS and QN

**Funding Acquisition**: TFS and QN

**Investigation**: YQ, LF, TFS and QN

**Methodology**: YQ, LF, TFS and QN

**Project Administration**: TFS and QN

**Resources**: TFS and QN

**Software**: YQ and QN

**Supervision**: TFS and QN

**Validation**: YQ, LF, TFS and QN

**Visualization**: YQ, LF, TFS and QN

**Writing – Original Draft Preparation**: YQ, LF, TFS and QN

**Writing – Review & Editing**: YQ, LF, TFS and QN

## Methods

### Ethics statement

All animal work performed in this study was approved by the University of California Irvine Institutional Animal Care and Use Committee (Protocol #AUP-20-145).

### Animal Husbandry

All animals used in this study were raised and handled in accordance with the guidelines of the Institutional Animal Care and Use Committee at University of California, Irvine. AB strain embryos were collected from natural crosses, raised at 28.5 °C in embryo medium (EM), and staged as previously described [79].

### Whole-mount in situ hybridization

In situ hybridization was performed on whole embryos as previously described [80]. Digoxygenin-and fluorescein-labelled riboprobes for *aldh1a2* [81], *krox20* [82], *otx2* [83] were synthesized using an RNA labelling kit (Roche) from cDNA that had been previously cloned into PCS2+ plasmids and linearized.

### Imaging and measurement of hindbrain

Embryos were flat mounted in glycerol as previously described [84] and imaged on a Zeiss Axioplan 2 compound microscope equipped with a Micropublisher 5.0 RTV camera with Zeiss ZEN 3.1 (blue edition) software. Hindbrain measurements were performed using ImageJ/Fiji software.

### Computational domains of the model

The entire hindbrain along with the RA production region, modeled as the “morphogen domain”, is used to model the diffusion and distribution of morphogens (Fig 1B). In the two-dimensional model, the morphogen domain is assumed as a rectangle with the anterior-posterior (A-P) axis as its length and the left-right (L-R) axis as its width. We take the posterior end of the MHB, defined by *otx2* expression, as the anterior limit of the domain, *x*^(1)^ = 0 and the posterior end of the RA production region, defined by *aldh1a2* expression, as its posterior limit, *x*^(1)^ = *L*_1_ (*t*). The L-R width of the hindbrain is *L* (*t*). The morphogen domain has a rectangular structure with dynamic sizes:

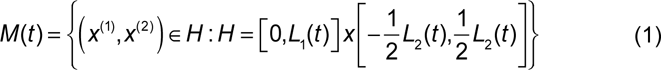

The RA production region is modeled as:

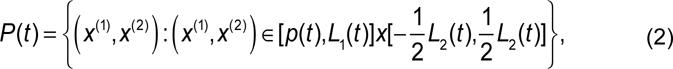

where *p*(*t*) is the A-P position of the anterior boundary of the RA production region at time *t*. *L*_1_(*t*), *L*_2_(*t*) and *p*(*t*) are obtained from experimental measurements made in zebrafish embryos at 11, 12, 13 and 14 hours postfertilization (hpf). A cubic interpolation is used to obtain the smooth curves (Fig 2A, B).

Individual cells are modeled in a “tissue domain” that is contained within the morphogen domain. The tissue domain shares the same L-R axis with of the morphogen domain and its A-P range is proportional to the range of morphogen:

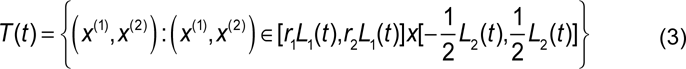

where *r*_1_ and *r*_2_ are constants given in the Table S1. At ***x*** = (*x*_1_,*x*_2_), the growth velocity of the tissue is given by

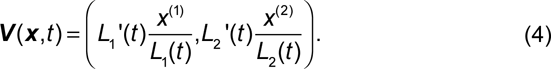

### Stochastic dynamics of morphogens

To model morphogen dynamics in the growing hindbrain, we use stochastic convection-reaction-diffusion equations. The equations for RA are given by

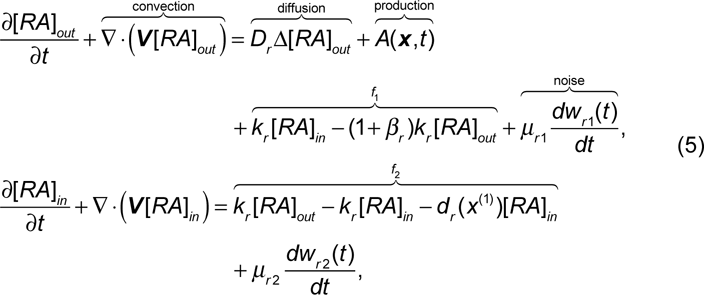

where [*RA*]*_out_* and [*RA*]*_in_* are extracellular and intracellular forms of RA, respectively. 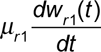 and 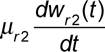 are additive white noise. The convection term describes the dilution and advection of RA caused by convergence extension. The production is confined to the RA production region and modeled by a Hill function of AP position *x*^(1)^ with a large Hill coefficient:

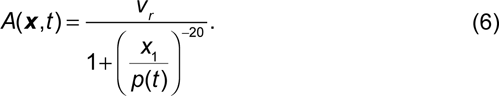

In *f_1_* and *f_2_*, *k_r_* is the rate of exchange of morphogen between intracellular and extracellular forms. The rate of extracellular morphogen degradation is taken as a constant *β_r_k_r_* and the degradation of intracellular morphogen rate *d_r_* is a piecewise function with

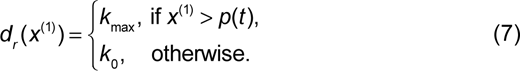

The degradation rate in the RA production region has a large value *k*_max_, since the RA degrading enzyme cyp26a1 is expressed in the RA production region [28]. We use an absorbed boundary condition of *x^(1)^*=0 since cyp26a1 is highly expressed in the forebrain and MHB, providing a sink for RA. No-flux boundary conditions are used for the other three boundaries.

Similarly, we model both free diffusible FGF (*[Fgf]_free_*) and FGF (*[Fgf]_signal_*) signaling as the following:

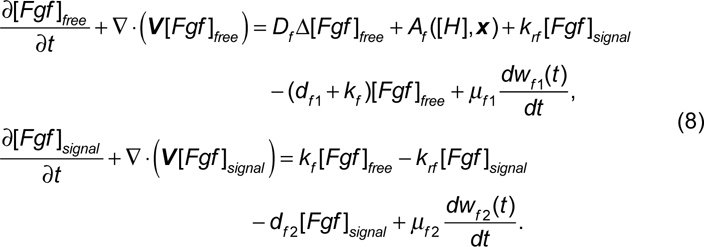

The free diffusible FGF binds with its receptor to form a complex with rate *k_f_[Fgf]_free_*. The complex between FGF and its receptor represents the FGF signal (*[Fgf]_signal_*) for simplification. The term *k_rf_[Fgf]_signal_* describes the dissociation rate of the complex. *d_f1_* and *d_f2_* are degradation rates of free diffusible FGF and FGF signaling, respectively. The production of FGF is upregulated by *hoxb1a* and the production rate is modelled by a Hill function for *hoxb1a*:

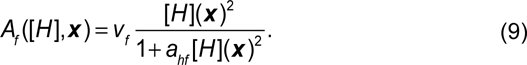

The *hoxb1a* level [*H*] is defined at the center of each cell. In Eq. (9), the term [*H*](***x***), defined at arbitrary location ***x***, is obtained by interpolating [*H*] values with locations in cell centers (**Supplementary Materials S1**). No-flux boundary conditions are used for FGF at all four boundaries.

### Stochastic dynamics of downstream genes

We model the dynamics of gene expression with a system of stochastic differential equations based on the gene network (Fig 1C). For the *i*-th cell centered at ***c****_i_*, the equations for the gene expression are given by

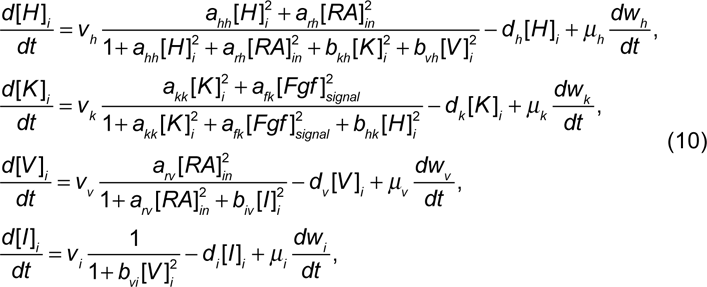

where [*H*], [*K*], [*V*] and [*I*] are gene expression of *hoxb1a*, krox20, *vhnf1* and *irx3*, respectively. 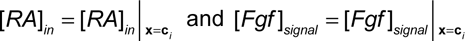 refer to the RA and FGF signaling levels at the center of i-th cell ***c****_i_* to provide spatial signals for cells.

### Models for individual cells and their interactions

Following our previous study [16], we use the subcellular element method (SCEM) to model individual cells [56]. A total of *N_cell_*=345 cells are modeled in each simulation, where 23 rows and 15 columns of cells align uniformly in the rectangular tissue region at the initial stage (11 hpf). Each cell consists of sub-cellular elements (nodes) and interacts according to a prescribed intercellular force potential. A cell consists of 2*N_node_* (*N_node_*=6) nodes and those nodes form two hexagonal layers (**Fig S6A)**. The radius of the outer layer is *R_out_* and the radius of the inner layer is *R_in_*. Initially cells are uniformly distributed in the tissue domain (Fig 4B). For a system with *N_cell_* cells and 2*N_node_* nodes per cell, the location of *i*-th node in *n*-th cell x*_n_*_,*i*_ is determined by the equation

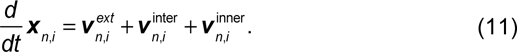

On the right hand side, the first term represents cell migration due to convergent-extension [85, 86]. It is given by

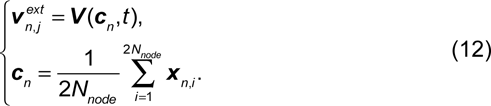

The second term represents the forces between cells while the third term represents forces between nodes within the same cell to maintain stable cell morphologies [56] (see **Supplementary Materials S2**).

### Definition of boundary location (*m*), sharpness index (*SI*) and number of dislocated cells (*DC*)

To study boundary locations and sharpness quantitatively, we define three quantities: boundary location (m), boundary sharpness index (SI) and number of dislocated cells (DCs). For example, the A-P position of the r3/r4 boundary is denoted by *m*(*r* 3 *r* 4) and its boundary SI is denoted by *SI*(*r* 3 *r* 4). A cell is called a DC if: a) its identity is different from the segment in which it is located; b) its distance to the boundary of its correct segment is over three cell-diameters.

In a region with A-P coordinates in the range of (*a*,*b*), we split the index set of all cells into two sets *S_L_* and *S_R_* based on cell identities, where cells in *S*_L_ or *S*_R_ have segmental identities located anterior or posterior to this region. We define the distance function from the *i*-th cell centered at ***c****_i_* to an arbitrary straight line with A-P position *k* (the potential location of boundary):

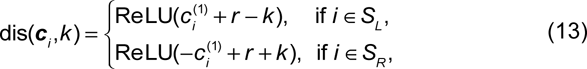

where *r* is the radius of the cell, and

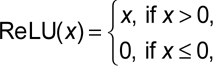

is the rectified linear unit function. For a cell in *S*_L_ or *S*_R_ with non-zero distance, this distance function calculates the Euclidean distance between the anterior or posterior distal ends of this cell to the potential location of boundary *k*. This distance function is illustrated in **Fig S6B**.

We quantify the boundary location (*m*), SI and number of DCs in this region, called *K*, by solving an optimization problem:

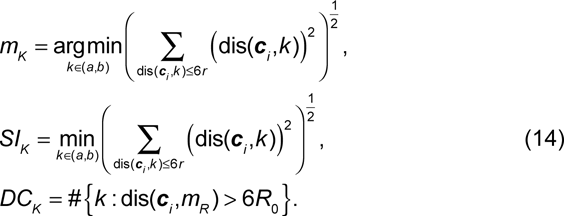

Particularly if the distance from a cell to the boundary is within three cell-diameters, the cell contributes to the boundary location and boundary SI. Otherwise, it is regarded as a DC and it does not contribute to the calculation of either boundary location or sharpness.

Next, we split all cells in the responding tissue domain with index set *S* into three sets with distinct cell types at time *t*. There are *hoxb1a* cells (*S*_h_), *krox20* cells (*S*_k_) and non-expressing cells (*S*_n_) based on their expression level of *hoxb1a* and *krox20*.

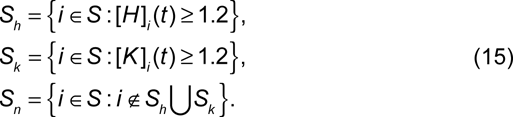

Now, we calculate those quantities for four boundaries in the tissue domain one-by-one by utilizing Eq. (14) and Eq. (15) as shown in the flow below:

#### Algorithm 1: Calculate m, SI and DC for cells in domain with AP range [*r*_1_*L*_1_(*t*),*r*_2_*L*_1_(*t*)]

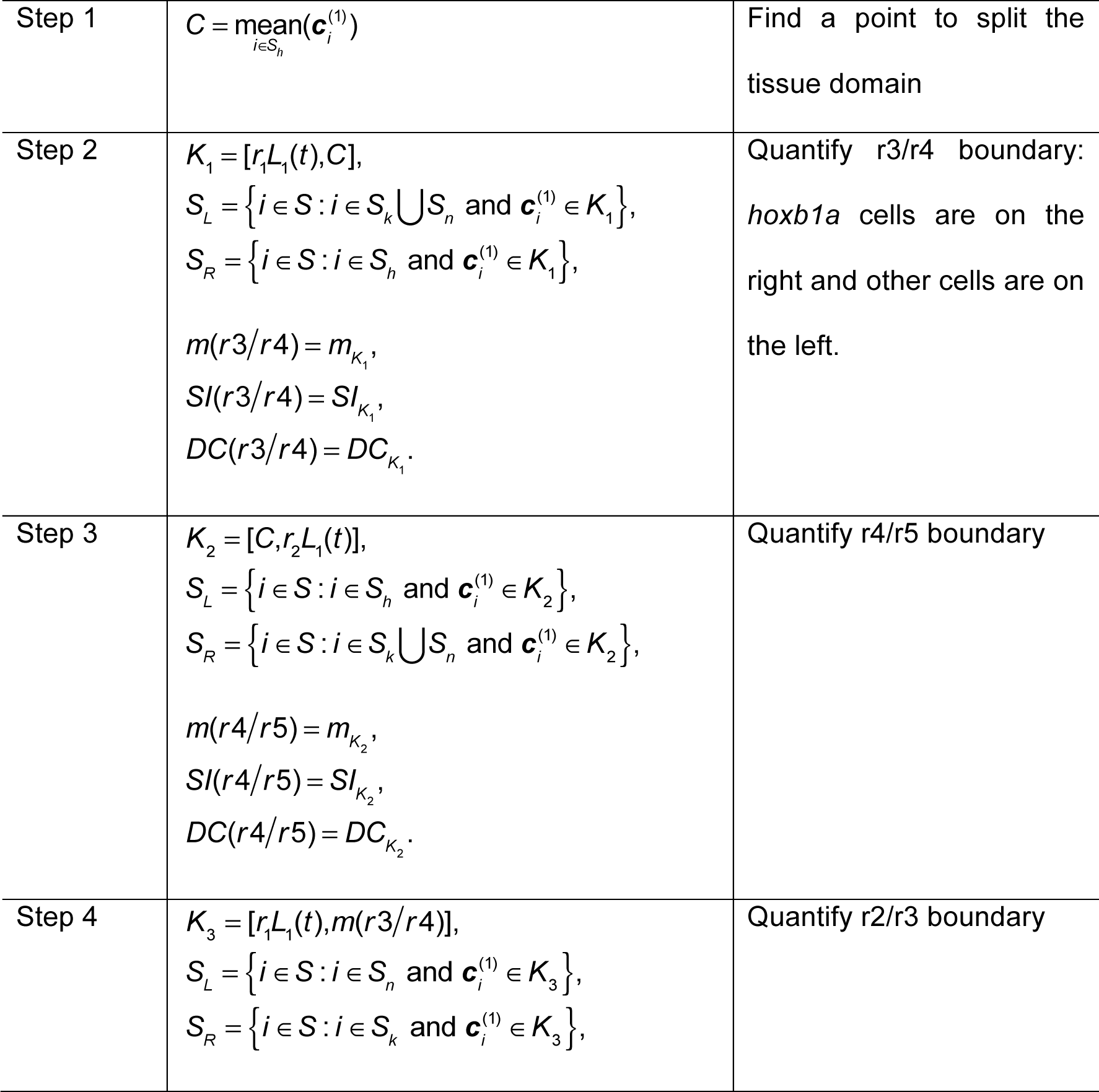
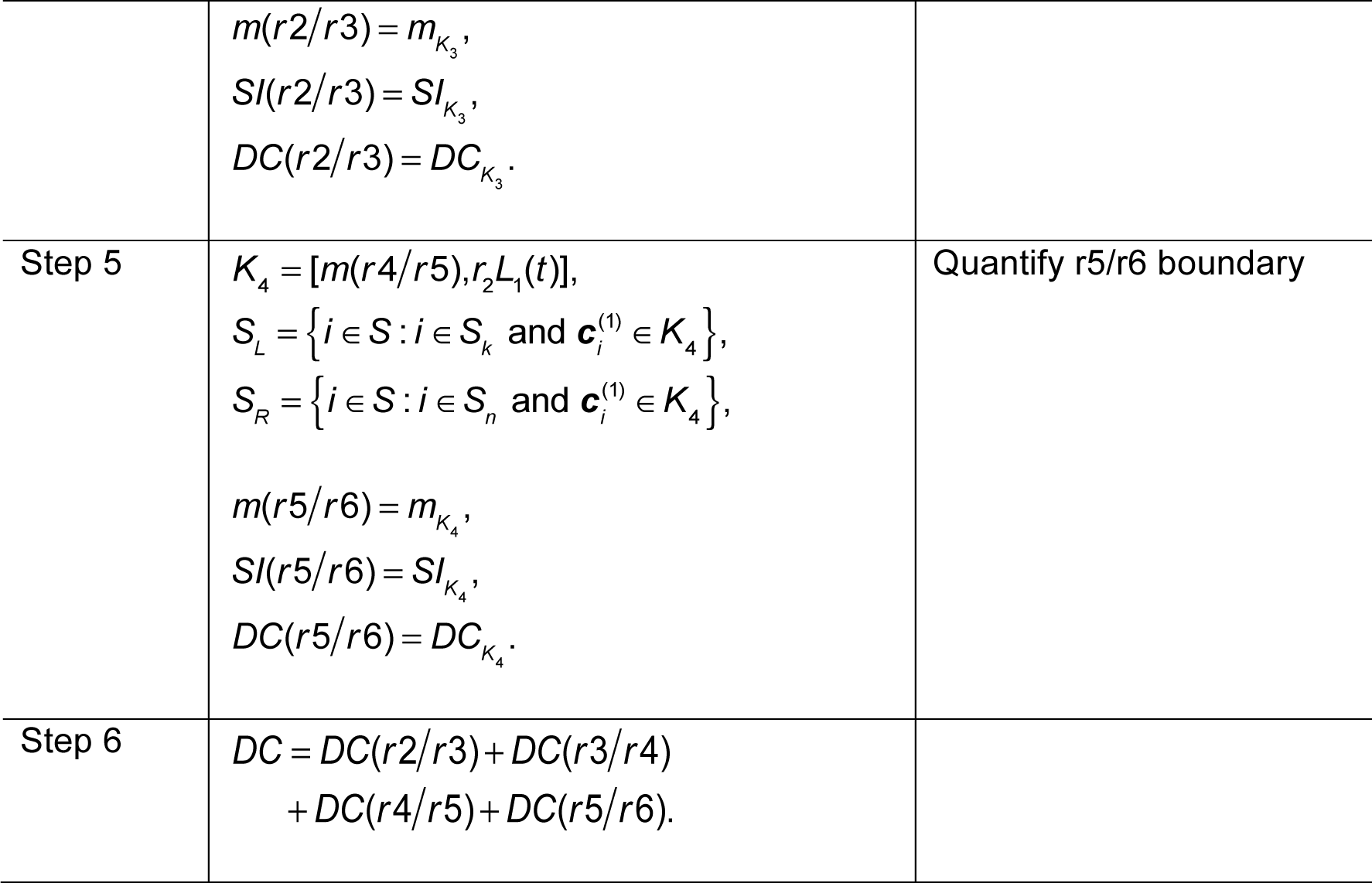

## Supporting information

**S1 Text. This file contains Supplementary Material including modeling details, parameter values, and experimental data.**

**S1 Figure. This file contains Supplementary Figures.**

