## Supplementary Information for "Multiple morphogens and rapid elongation promote segmental patterning during development"

1. Department of Mathematics, University of California, Irvine, CA 92697, USA
2. Current Address: Department of Mathematics, Michigan State University, MI 48824, USA
3. Department of Developmental and Cell Biology, University of California, Irvine, CA 92697, USA
4. The NSF-Simons Center for Multiscale Cell Fate Research, University of California, Irvine, CA 92697, USA

### S1. Signaling exchanges between continuum domain and individual cells

In the computational framework, the morphogen domain uses a regular rectangular mesh and the tissue domain uses an irregular mesh, where grids are the nodes in each moving cell. Morphogens need to transmit signal to individual cells and FGF synthesis needs *hoxb1a* expression in individual cells. We use interpolations to conduct the signaling exchanges from two domains.

From morphogen domain to tissue domain, we use a constant interpolation. For $i$-th cell, the morphogen level $[M]_i$ that is closest to the cell location:

$$\begin{cases} [M]_i = [M]\Big|_{\boldsymbol{x}=\left(x_1^{i_0}, x_2^{j_0}\right)}, \\ \left(i_0, j_0\right) = \underset{i,j}{\arg\min}\left|\left(x_1^i, x_2^j\right) - \boldsymbol{c}_i\right|, \end{cases} \tag{S1}$$

Here $[M]_i$ can represent either the intracellular RA ($[RA]_{in}$) or FGF signaling ($[Fgf]_{signal}$) in *i*-th cell. From tissue domain to morphogen domain, we use the build-in function "*griddata*" in MATLAB to interpolate the *hoxb1a* expression [*H*] defined on scatter points (i.e. centers of individual cells) to the regular mesh in the morphogen domain:

$$[H](\boldsymbol{x}) = griddata([H]_i\Big|_{i=1}^{N_{cell}}, \boldsymbol{c}_i\Big|_{i=1}^{N_{cell}}, \boldsymbol{x}) \tag{S2}$$

**S2. Models for individual cells and their interactions**

Following our previous work [1], we use the subcellular element method (SCEM) to model individual cells [2]. Each cell consists of sub-cellular elements (nodes) and interacts according to a prescribed intercellular force potential. A cell consists of $2N_{node}$ ($N_{node}$=6) nodes and those nodes form two hexagonal layers as shown in **Fig S6A**. The radius of the outer layer is $R_{out}$ and the radius of the inner layer is $R_{in}$. Initially cells are uniformly distributed in the tissue domain as shown in **Fig 2H**. For a system with $N_{cell}$ cells and each cell has $2N_{node}$ nodes, the location of *i*-th node in *n*-th cell $\boldsymbol{x}_{n,j}$ is determined by the equation

$$\frac{d}{dt}\boldsymbol{x}_{n,i} = \boldsymbol{v}_{n,i}^{\text{ext}} + \boldsymbol{v}_{n,i}^{\text{inter}} + \boldsymbol{v}_{n,i}^{\text{inner}}. \tag{S3}$$

On the right hand side, the first term represents cell migration due to convergent extension [3, 4]. It is given by

$$\begin{cases} \boldsymbol{v}_{n,j}^{ext} = \boldsymbol{V}(\boldsymbol{c}_n, t), \\ \boldsymbol{c}_n = \dfrac{1}{2N_{node}} \displaystyle\sum_{i=1}^{2N_{node}} \boldsymbol{x}_{n,i}. \end{cases} \tag{S4}$$

The second term represents the forces between cells while the third term represents forces between nodes within the same cell to maintain the stable cell morphology [2]. For two nodes, their interactions are determined by Morse potential with a cutoff distance $d$:

$$\Phi(r)=\begin{cases} U\exp(-\frac{r}{\xi})-V\exp(-\frac{r}{\zeta}), & \text{if } r\le d; \\ 0, & \text{otherwise}; \end{cases} \tag{S5}$$

where $r$ is the distance between these two nodes and $d$ is the constant cut-off distance taken as two-fold of cell diameter ($4R_0$=16 μm). The short-range repulsion $U\exp(-\frac{r}{\xi})$ maintains a minimum distance between nodes and the long-range attraction $-V\exp(-\frac{r}{\zeta})$ makes nodes move closer. For nodes in the same cell, the *non-zero* intracellular forces are determined by:

$$\Phi_{\text{intra}}(|\boldsymbol{x}_{n,i}-\boldsymbol{x}_{n,j}|)=U_{\text{intra}}\exp\left(-\frac{|\boldsymbol{x}_{n,i}-\boldsymbol{x}_{n,j}|}{\xi_{\text{intra}}}\right)-V_{\text{intra}}\exp\left(-\frac{|\boldsymbol{x}_{n,i}-\boldsymbol{x}_{n,j}|}{\zeta_{\text{intra}}}\right) \tag{S6}$$

To model selective cell sorting for nodes in two cells, the *non-zero* intercellular forces depend on the identities of these two cells. In some cases, cells with the same identity sort using both long-range attractive and short-range repulsive forces, and cells with different identities sort using repulsive forces:

$$\Phi_{\text{inter}}(r)=\begin{cases} U_{\text{inter}}^{\text{Atr}}\exp(-\frac{r}{\xi_{\text{inter}}})-V_{\text{inter}}^{\text{Atr}}\exp(-\frac{r}{\zeta_{\text{inter}}}), & \text{same identity;} \\ U_{\text{inter}}^{\text{Rep}}\exp(-\frac{r}{\xi_{\text{inter}}}), & \text{different identities.} \end{cases} \tag{S7}$$

In our model, selective inter-cellular forces depend on *krox20* levels in a pair of cells. We determine the inter-cellular forces using an affine combination of two types of forces in Eq. (S7). The discrepancy between *krox20* levels in these cells is used to determine the coefficients of the affine combination. For $m$-th and $n$-th cell, a similarity function is defined to describe such a discrepancy:

$$\mathcal{F}(m,n) = \frac{1}{2}(\delta^m \delta^n + 1) \in [0,1], \tag{S8}$$

where $\delta^i$ is the relative level of *krox20* in $i$-th cell normalized by the maximum expression *krox20* over all cells. $\delta^i = 1$ indicates that the cell expresses a maximum level of *krox20* and $\delta^i = -1$ indicates the cell has zero *krox20* expression. The force between two cells is attractive when they have similar *krox20* levels ($\mathcal{F} = 1$ if $\delta_m = \delta_n = \pm 1$) and is repulsive when they have different *krox20* levels ($\mathcal{F} = 0$ if $\delta_m = -\delta_n = \pm 1$). The Morse potentials between a pair of nodes in two different cells with locations at $\boldsymbol{x}_{m,j}$ and $\boldsymbol{x}_{n,i}$ are defined by

$$\begin{aligned}\Phi_{\text{inter}}(|\boldsymbol{x}_{n,i} - \boldsymbol{x}_{m,j}|) = &\left[\left(1 - \mathcal{F}(m,n)\right)U_{\text{inter}}^{\text{Rep}} + \mathcal{F}U_{\text{inter}}^{\text{Atr}}\right]\exp\left(-\frac{|\boldsymbol{x}_{n,i} - \boldsymbol{x}_{m,j}|}{\xi_{\text{inter}}}\right) \\ &- \mathcal{F}(m,n)\mathrm{V}_{\text{inter}}^{\text{Atr}}\exp\left(-\frac{|\boldsymbol{x}_{n,i} - \boldsymbol{x}_{m,j}|}{\zeta_{\text{inter}}}\right).\end{aligned} \tag{S9}$$

Taken together, the second term in Eq. (S3) given by

$$\boldsymbol{v}_{n,i}^{\text{inter}} = \eta_{n,i} - \nabla_{\boldsymbol{x}_{n,i}}\left[\sum_{j \neq i}^{2N_{node}} \Phi_{\text{intra}}(|\boldsymbol{x}_{n,i} - \boldsymbol{x}_{n,j}|) + \sum_{m \neq n}^{N_{cell}} \sum_{j}^{2N_{node}} \Phi_{\text{inter}}(|\boldsymbol{x}_{n,i} - \boldsymbol{x}_{m,j}|)\right], \tag{S10}$$

where $\eta_{n,i} \sim \mathcal{N}(0,10)$ is the Gaussian-distributed noise.

The canonical SCEM only considers Morse potentials as discussed above. To avoid squashed or exploded cells, we add other spring-type pairwise interactions to nodes within the same cell [1]. In $i$-th cell, we evenly divide nodes into two layers, where each layer initially has a regular polygonal structure. These pairwise interactions act on neighboring nodes between the same layer $\boldsymbol{x}_{n,i} \sim \boldsymbol{x}_{n,i\pm 1}$ and the nodes between different layers with the same index $\boldsymbol{x}_{n,i} \sim \boldsymbol{x}_{n,i\pm N_{node}}$. The force is given by

$$\Psi(\boldsymbol{x}_{n,i}, \boldsymbol{x}_{n,j}) = \mu \frac{|\boldsymbol{x}_{n,i} - \boldsymbol{x}_{n,j}| - l^{n}_{i,j}}{|\boldsymbol{x}_{n,i} - \boldsymbol{x}_{n,j}|} (\boldsymbol{x}_{n,i} - \boldsymbol{x}_{n,j}), \tag{S11}$$

where $l^{n}_{i,j}$ is the constant length between node $\boldsymbol{x}_{n,i}$ and $\boldsymbol{x}_{n,j}$ at an initial time and it has three possible values, $l_{out}$, $l_{in}$, $l_{inter}$, depending on the types of two nodes (**Fig S6A**). Then for *i*-th node in the *n*-th cell, the motion from this force is

$$\boldsymbol{v}^{\text{inner}}_{n,i} = \Psi(\boldsymbol{x}_{n,i}, \boldsymbol{x}_{n,i+1}) + \Psi(\boldsymbol{x}_{n,i}, \boldsymbol{x}_{n,i-1}) + \Psi(\boldsymbol{x}_{n,i}, \boldsymbol{x}_{n,i\pm N_{node}}). \tag{S12}$$

To prevent cells leaving the tissue domain, we add two layers of “ghost cells” located outside the boundary tissue domain to provide extra forces.

### S3. One-dimensional gene expression model

In the main text, we compare and contrast the two-morphogen and one-morphogen models. The equations are defined for the fixed one-dimensional domain with deterministic description. The equations for the two-morphogen model are given by Eq. (10) in Methods by removing noise terms. For the one-morphogen model, RA induces both *hoxb1a* and *krox20*. Only equations for *hoxb1a* and *krox20* are modified:

$$\begin{aligned} \frac{d[H]}{dt} &= v_h \frac{a_{hh}[H]^2 + a_{rh}[RA]^2_{in}}{1 + a_{hh}[H]^2 + a_{rh}[RA]^2_{in} + b_{kh}[K]^2 + b_{vh}[V]^2} - d_h[H] \ , \\ \frac{d[K]}{dt} &= v_k \frac{a_{kk}[K]^2 + a_{rk}[RA]^2_{in}}{1 + a_{kk}[K]^2 + a_{rk}[RA]^2_{in} + b_{hk}[H]^2} - d_k[K] \ . \end{aligned} \tag{S13}$$

**S4. Initial cell distribution for cell sorting-only model.**

In **Fig 5B**, we use mixture Gaussian distribution to generate the "salt-and-pepper" initial cell distribution. We label the cells with r2, r3, r4, r5 and r6 identities as 1, 2, 3, 4 and 5, respectively. For a cell with A-P position x, we normalized its A-P position $\tilde{x} = \frac{x - x_{\min}}{x_{\max} - x_{\min}}$ in the range of [0,1] where $x_{\min} = r_1 L_1(0)$ and $x_{\max} = r_2 L_1(0)$ are A-P positions of boundaries of the tissue domain. The probability for this cell being *i*-th type of cell is given by

$$p_i(\tilde{x}) = \frac{\phi_i \mathcal{N}(\tilde{x} | \mu_i, \sigma_i)}{\sum_{i=1}^{5} \phi_i \mathcal{N}(\tilde{x} | \mu_i, \sigma_i)}. \quad \text{(S14)}$$

We take the weights and the variance as $\phi_i = 1/5$ $\sigma_i = 0.08$ for all *i*. The means $(\mu_1, \mu_2, \mu_3, \mu_4, \mu_5) = \frac{1}{10}(1,3,5,7,9)$ denote the average A-P positions of each cell type. The probability distributions of different cell types are shown in **Fig S6C**.

**S5. Determination of cell fate**

In the main text, we discuss cell commitment time, defined by the latest time that a cell changes its fate. To determine cell fate numerically, we used thresholds for *hoxb1a* and *krox20* expression. There are three types of cells in the r2-r6 domain. Cells in r2 and r6 have low expression of both *hoxb1a* and *krox20*. Cells in r3 and r5 have low *hoxb1a* and high *krox20* expression. Cells in r4 have high *hoxb1a* and low *krox20* expression. "High" and "low" levels are defined by expression thresholds of 1.2, with no cell identified as having high expression of both *hoxb1a* and *krox20*. The expression thresholds are defined as the following:

1) none expressing identity: $[H] < 1.2$, $[K] < 1.2$;
2) *hoxb1a* identity: $[H] < 1.2$, $[K] \geq 1.2$; (S15)
3) *krox20* identity: $[H] \geq 1.2$, $[K] < 1.2$.

### S6. Numerical solvers

#### S6.1. Solving PDEs (morphogens) in moving boundaries

For morphogens RA and FGF, the two-dimensional spatial domains have moving boundaries. A general form of the equation is given by

$$\begin{aligned}&\frac{\partial[M]}{\partial t} + \nabla\cdot\left(\boldsymbol{V}[M]\right) = D\Delta[M] + f,\\&\boldsymbol{x} = (x^{(1)}, x^{(2)}) \in [0, L_1(t)]x[-\frac{1}{2}L_2(t), \frac{1}{2}L_2(t)].\end{aligned} \tag{S16}$$

To numerically solve these PDEs, we first transform the dynamic domain to the unit square by using a coordinate transformation:

$$\begin{cases}\boldsymbol{X} = (X^{(1)}, X^{(2)}) = \left(\frac{1}{L_1(t)}x^{(1)}, \frac{1}{L_2(t)}x^{(2)} + \frac{1}{2}\right) \in [0,1]x[0,1],\\ \tau = t.\end{cases} \tag{S17}$$

The partial derivatives in a transformed coordinate system are given by

$$\begin{aligned}&\nabla_{\boldsymbol{X}} = \left(\frac{\partial}{\partial X^{(1)}}, \frac{\partial}{\partial X^{(2)}}\right) = \left(L_1(t)\frac{\partial}{\partial x^{(1)}}, L_2(t)\frac{\partial}{\partial x^{(2)}}\right),\\&\frac{\partial^2}{\partial\left(X^{(1)}\right)^2} = \left(L_1(t)\right)^2\frac{\partial^2}{\partial\left(x^{(1)}\right)^2},\\&\frac{\partial^2}{\partial\left(X^{(2)}\right)^2} = \left(L_2(t)\right)^2\frac{\partial^2}{\partial\left(x^{(2)}\right)^2},\\&\frac{\partial}{\partial\tau} = \frac{\partial}{\partial t} + L_1^{'}(t)X^{(1)}\frac{\partial}{\partial x^{(1)}} + L_2^{'}(t)\left(X^{(2)} - \frac{1}{2}\right)\frac{\partial}{\partial x^{(2)}}.\end{aligned} \tag{S18}$$

The resulting equation in the new coordinate system is given by:

$$\frac{\partial [M]}{\partial \tau} = \frac{1}{\left(L_1(t)\right)^2}\frac{\partial^2 [M]}{\partial \left(X^{(1)}\right)^2} + \frac{1}{\left(L_2(t)\right)^2}\frac{\partial^2 [M]}{\partial \left(X^{(2)}\right)^2} + \frac{L_1'(t)}{L_1(t)}[M] + \frac{L_2'(t)}{L_2(t)}[M] + f \quad \text{(S19)}$$

For spatial discretization, we use uniform rectangular grids with 151 and 71 grid points along the A-P and L-R axis, respectively. The central difference method is used for the discretization of the diffusion term. For temporal discretization, we use the Euler-Maruyama method with a fixed time step. The time step for morphogens and gene expressions model is $\Delta t_1 = 1.8$ seconds.

#### S6.2. Solving equations in discrete cells model

We use the Euler-Maruyama method to solve Eq. (S3) with the fixed time step $\Delta t_2 = 3.6$ seconds. For calculations of the pairwise interactions between nodes in Eq. (S10), we use GPUs algorithm by using "gpuArray()" function in MATLAB.

### S7. Parameters

#### S7.1. Parameters value and selection

Parameter values used in our simulations are listed in **Tables S1, S2** and **S3**. In particular, parameters used in cell mechanical models are listed in **Table S1**, and parameters used in gene regulation models are listed in **Table S2**. The sampling ranges of random parameters in **Fig 7** are listed in **Table S3**.

In cell-based models, the A-P range of the tissue domain, determined by ratios $r_1$ and $r_2$, is chosen to allow r2-r6 to be contained within this region, where r3-r5 A-P lengths are from experimental measurement (**Table S7**) while r2 and r6 lengths are assumed to have similar lengths to r3 and r5 (since all rhombomeres in r2-r6 have similar A-P lengths). The cell radius, $R_{\text{out}}$, is around 4 μm in 11-14hpf, which is estimated from images in [5]. In cell mechanical models, the parameters are taken from our previous work [1] and the time and length units have been adjusted based on new measurements. In gene regulation models, the parameters for RA (i.e. $D_r$, $v_r$, $\beta_r$, $k_{max}$, and $k_0$) are taken from previous modeling and experimental studies [6]. Since FGF generated in r4 is likely to have a shorter decay length than RA, we use a 3-fold smaller diffusion coefficient $D_f$ than for RA ($D_r$). The degradation rate chosen, $d_{f2}$, is also larger than the degradation rate of RA. As a result, FGF gradients are established much faster than RA gradients, Parameters for *hoxb1a* and *krox20* expression, $a_{hh}$, $a_{kk}$, $a_{rh}$, $d_h$, $d_k$ $b_{hk}$ and $b_{kh}$ were initially taken from our one-morphogen model [7] and adjusted manually to achieve values that produce a five-segment pattern according to length and time measured in this work.. All other parameters in the gene regulation models were adjusted to produce a five-segment pattern with correct rhombomere A-P lengths. The noise magnitudes, ($\mu_*$ parameters) were selected manually to give effective boundary sharpening that changed with increased or decreased noise magnitudes.

| **Parameters** | **Value** | **Unit** | **Reference** |
|---|---|---|---|
| $r_1$ | 0.038 | -- | -- |
| $r_2$ | 0.28 | -- | -- |
| $N_{column}$ | 15 | -- | -- |
| $N_{row}$ | 23 | -- | -- |
| $N_{cell}$ | 345 | -- | -- |
| $N_{node}$ | 6 | -- | -- |
| $R_{out}$ (cell radius) | 3.60 | μm | Estimated from images in [5] |

| | | | |
|---|---|---|---|
| $R_{in}$ | 1.80 | μm | [1] |
| $l_{out}$ | 3.60 | μm | |
| $l_{in}$ | 1.80 | μm | |
| $l_{inter}$ | 1.80 | μm | |
| $\mu$ | 1.11 | $sec^{-1}$ | |
| $U_{\text{intra}}$ | $1.33x10^{-1}$ | $μm^2\ sec^{-1}$ | |
| $\xi_{\text{intra}}$ | 2.4 | μm | |
| $V_{\text{intra}}$ | $5.56x10^{-2}$ | $μm^2\ sec^{-1}$ | |
| $\zeta_{\text{intra}}$ | 3.6 | μm | |
| $U_{\text{inter}}^{\text{Atr}}$ | $1.04x10^{-1}$ | $μm^2\ sec^{-1}$ | |
| $U_{\text{inter}}^{\text{Rep}}$ | $2.50x10^{-2}$ | $μm^2\ sec^{-1}$ | |
| $\xi_{\text{inter}}$ | 5.0 | μm | |
| $V_{\text{inter}}^{\text{Atr}}$ | $6.67x10^{-2}$ | $μm^2\ sec^{-1}$ | |
| $\zeta_{\text{inter}}$ | 11.00 | μm | |

**Table S1. Parameters for the cell mechanical model.**

| **Parameters** | **Value** | **Unit** | **Reference** |
|---|---|---|---|
| $D_r$ | 2.83 | $μm^2\ sec^{-1}$ | [6] |
| $v_r$ | $1.11x10^{-2}$ | $sec^{-1}$ | |
| $k_r$ | $2.22x10^{-4}$ | $sec^{-1}$ | |
| $\beta_r$ | 1 | -- | |
| $k_{\max}$ | $5.56x10^{-1}$ | $sec^{-1}$ | |

| | | | |
|---|---|---|---|
| $k_0$ | $1.11 \times 10^{-4}$ | $sec^{-1}$ | |
| $D_f$ | 0.85 | $\mu m^2\ sec^{-1}$ | -- |
| $v_f$ | $5.56 \times 10^{-2}$ | $sec^{-1}$ | -- |
| $a_{hf}$ | 2 | -- | -- |
| $k_f$ | $2.67 \times 10^{-4}$ | $sec^{-1}$ | -- |
| $k_{rf}$ | $2.67 \times 10^{-4}$ | $sec^{-1}$ | -- |
| $d_{f1}$ | $2.67 \times 10^{-4}$ | $sec^{-1}$ | -- |
| $d_{f2}$ | 0.013 | $sec^{-1}$ | -- |
| $\mu_{r1}$ | 0.1 | -- | -- |
| $\mu_{r2}$ | 0.003 | -- | -- |
| $\mu_{f1}$ | 0.1 | -- | -- |
| $\mu_{f2}$ | 0.01 | -- | -- |
| $v_h$ | 0.056 | $sec^{-1}$ | -- |
| $v_k$ | 0.056 | $sec^{-1}$ | -- |
| $v_v$ | 0.11 | $sec^{-1}$ | -- |
| $v_i$ | $3.33 \times 10^{-4}$ | $sec^{-1}$ | -- |
| $a_{hh}$ | 0.85 | -- | -- |
| $a_{rh}$ | 0.13 | -- | -- |
| $a_{kk}$ | 0.9 | -- | -- |
| $a_{fk}$ | 6 | -- | -- |
| $a_{rv}$ | 0.1 | -- | -- |
| $b_{kh}$ | 40 | -- | -- |
| $b_{vh}$ | 5 | -- | -- |

| | | | |
|---|---|---|---|
| $b_{hk}$ | 20 | -- | -- |
| $b_{iv}$ | 3.5 | -- | -- |
| $b_{vi}$ | 5 | -- | -- |
| $d_h$ | 0.022 | $sec^{-1}$ | -- |
| $d_k$ | 0.022 | $sec^{-1}$ | -- |
| $d_v$ | 0.022 | $sec^{-1}$ | -- |
| $d_i$ | $3.33 \times 10^{-4}$ | $sec^{-1}$ | -- |
| $\mu_h$ | 0.01 | -- | -- |
| $\mu_k$ | 0.01 | -- | -- |
| $\mu_v$ | 0.005 | -- | -- |
| $\mu_i$ | 0.005 | -- | -- |

**Table S2. Parameters for the equations of morphogens and intracellular genes.**

#### S7.2 Parameters sensitivity analysis

##### S7.2.1 Random parameters perturbations

In **Fig 7** and **Fig S5**, we generate the simulations with random parameters for the equations for *hoxb1a* and *krox20*. We use Latin hypercube sampling to generate high dimension random numbers and each sample is written as $(\omega_1,...,\omega_N)$. We list the parameters that are perturbed by using the random number. If not listed, those parameters are taken as the same as that in **Table S2**.

| **Parameters** | **Value** | **Unit** |
|---|---|---|
| $a_{hh}$ | $0.85x2^{-0.4+0.8\omega_1}$ | -- |
| $a_{kk}$ | $0.9x2^{-0.4+0.8\omega_2}$ | -- |
| $a_{rh}$ | $0.13x2^{-1+2\omega_3}$ | -- |

| | | |
|---|---|---|
| $a_{fk}$ | $6x2^{-1+2\omega_4}$ | -- |
| $b_{kh}$ | $40x2^{-1+2\omega_5}$ | -- |
| $b_{vh}$ | $5x2^{-1+2\omega_6}$ | -- |
| $b_{hk}$ | $20x2^{-1+2\omega_7}$ | -- |
| $d_h$ | $0.022x2^{-1+2\omega_8}$ | $sec^{-1}$ |
| $d_k$ | $0.022x2^{-1+2\omega_9}$ | $sec^{-1}$ |
| $b_{iv}$ | 4 | -- |
| $v_h$ | 2.5x$d_h$ | $sec^{-1}$ |
| $v_k$ | 2.5x$d_k$ | $sec^{-1}$ |

**Table S3. Parameters for Fig 7 and Fig S5.** If not specified, they are the same as that in Table S2.

**S7.2.2 Local sensitivity analysis of single parameters in gene regulation**

We performed single parameter perturbations to examine sensitivity to each parameter in our morphogen equations (Eq. (5-9)) and gene regulation equations (Eq. (10)). We systematically varied each parameter up and down by 20% of its total value and measured the effect on quantities including lengths of three rhombomeres, SIs of four boundaries and DC numbers. Specifically, we calculated the ratio between quantities in perturbed cases and quantities in unperturbed cases at the end of the simulations (14 hpf) (**Table S4**). The system is relatively insensitive to most parameters. It is relatively sensitive to $v_r$, $k_r$, $d_{f1}$ and $d_{f2}$ in morphogen equations. The robustness in morphogen dynamics can be improved by including many other mechanisms [8], which were not included in this work for simplification. In gene regulation models, the system is sensitive to auto-activation rate ($a_{hh}$), as also reported in previous work [1]. Since *hoxb1a* and *krox20* compete with each other, sensitivity to $v_h$, $v_k$, $d_h$ and $d_k$ are expected (many simulations fail to form rhombomeres) since they risk maximal values of *hoxb1a* or *krox20*. Alternatively, we can avoid these perturbations on maximal values of *hoxb1a* or *krox20* by coupling the ratio $v_h/d_h$ or $v_k/d_k$. The system turns out be robust to the perturbations on $v_h$ or $v_k$ with such coupling (**Table S4**).

| Parameter | Fold changed | Ratio of perturbed quantities to unperturbed quantities | | | | | | | | # failed simulations |
|---|---|---|---|---|---|---|---|---|---|---|
| | | r3 | r4 | r5 | SI(r2/r3) | SI(r3/r4) | SI(r4/r5) | SI(r5/r6) | DCs | |
| $D_r$ | 0.8 | 0.96 | 1.05 | 0.85 | 1.23 | 1.46 | 1.11 | 1 | 1.45 | 0 |
| | 1.2 | 1.02 | 0.9 | 1.09 | 0.93 | 0.94 | 1.37 | 1.14 | 2.26 | 0 |
| $v_r$ | 0.8 | 1 | 1.11 | 0.94 | 0.99 | 0.91 | 0.97 | 1.11 | 0.56 | 0 |
| | 1.2 | 0.95 | 0.92 | 1.07 | 1.44 | 1.83 | 1.39 | 0.98 | 4.39 | 1 |
| $\beta_r$ | 0.8 | 0.97 | 0.99 | 1.01 | 0.83 | 0.77 | 0.86 | 0.97 | 0.97 | 0 |
| | 1.2 | 0.97 | 0.99 | 1.01 | 0.83 | 0.77 | 0.86 | 0.97 | 0.97 | 0 |
| $k_{max}$ | 0.8 | 0.97 | 0.99 | 1.00 | 0.82 | 0.76 | 0.97 | 0.99 | 0.81 | 0 |
| | 1.2 | 0.97 | 0.98 | 1.02 | 0.82 | 0.76 | 0.90 | 0.97 | 0.97 | 0 |
| $k_r$ | 0.8 | -- | -- | -- | -- | -- | -- | -- | -- | 10 |
| | 1.2 | 0.92 | 1.18 | 0.84 | 1.1 | 1.66 | 2.31 | 0.82 | 4.84 | 1 |
| $D_f$ | 0.8 | 0.92 | 0.98 | 1.02 | 0.82 | 0.73 | 0.95 | 0.94 | 1.13 | 0 |
| | 1.2 | 1.02 | 1.03 | 0.95 | 0.9 | 0.7 | 1.23 | 1.33 | 0.89 | 0 |
| $k_f$ | 0.8 | 1.04 | 1.03 | 1.03 | 0.93 | 0.66 | 0.89 | 1.04 | 0.48 | 0 |
| | 1.2 | 0.9 | 0.97 | 0.95 | 0.66 | 0.91 | 1.04 | 1.16 | 1.61 | 0 |
| $d_{f1}$ | 0.8 | 1.12 | 0.95 | 1.18 | 0.85 | 0.75 | 0.64 | 1.07 | 0.4 | 0 |
| | 1.2 | 0.84 | 1.02 | 0.88 | 0.87 | 1.01 | 1.26 | 1.17 | 3.39 | 0 |
| $d_{f2}$ | 0.8 | 1.27 | 0.89 | 1.3 | 0.99 | 0.69 | 0.78 | 0.91 | 1.05 | 0 |

| | | | | | | | | | | |
|---|---|---|---|---|---|---|---|---|---|---|
| | 1.2 | 0.76 | 1.21 | 0.7 | 1.03 | 3.26 | 1.75 | 1.76 | 4.39 | 1 |
| $v_h$ | 0.8 | -- | -- | -- | -- | -- | -- | -- | -- | 10 |
| | 1.2 | -- | -- | -- | -- | -- | -- | -- | -- | 10 |
| $v_k$ | 0.8 | -- | -- | -- | -- | -- | -- | -- | -- | 10 |
| | 1.2 | 1.31 | 0.98 | 1.36 | 1.57 | 0.82 | 0.66 | 0.96 | 2.42 | 0 |
| $v_v$ | 0.8 | 1.31 | 1.39 | 1.14 | 0.89 | 0.61 | 0.94 | 0.72 | 0.4 | 0 |
| | 1.2 | 0.78 | 0.95 | 0.63 | 1.54 | 4.73 | 1.66 | 1.69 | 6 | 1 |
| $v_i$ | 0.8 | -- | -- | -- | -- | -- | -- | -- | -- | 10 |
| | 1.2 | 1.2 | 1.34 | 1.12 | 0.7 | 0.67 | 0.64 | 0.78 | 0.4 | 0 |
| $a_{hh}$ | 0.8 | 0.66 | 0.94 | 0.63 | 1 | 1.57 | 2.68 | 1.87 | 2.26 | 0 |
| | 1.2 | -- | -- | -- | -- | -- | -- | -- | -- | 10 |
| $a_{kk}$ | 0.8 | 0.75 | 0.97 | 0.83 | 0.7 | 0.93 | 1.2 | 1.4 | 3.39 | 0 |
| | 1.2 | 1.16 | 1 | 1.18 | 1.06 | 0.73 | 0.89 | 1.09 | 1.21 | 0 |
| $a_{rh}$ | 0.8 | 0.85 | 0.89 | 0.67 | 1.3 | 4.44 | 1.85 | 1.23 | 4.84 | 5 |
| | 1.2 | 1.26 | 1.22 | 1.21 | 1.19 | 1.12 | 0.92 | 0.99 | 0.81 | 0 |
| $a_{rv}$ | 0.8 | 1.29 | 1.36 | 1.19 | 0.95 | 0.59 | 1 | 0.62 | 0.4 | 0 |
| | 1.2 | 0.8 | 0.97 | 0.64 | 2.01 | 5.1 | 1.41 | 1.72 | 6.05 | 2 |
| $a_{fk}$ | 0.8 | 0.86 | 1.07 | 0.81 | 0.54 | 0.86 | 1.29 | 1.36 | 3.32 | 1 |
| | 1.2 | 1.09 | 0.93 | 1.13 | 0.86 | 0.85 | 0.84 | 1.11 | 0.73 | 0 |
| $b_{hk}$ | 0.8 | 0.94 | 0.95 | 1.05 | 0.68 | 0.74 | 0.71 | 1.08 | 0.89 | 0 |
| | 1.2 | 0.97 | 1.03 | 0.93 | 0.93 | 0.69 | 1.03 | 1.19 | 1.13 | 0 |
| $b_{kh}$ | 0.8 | 0.99 | 1.04 | 0.96 | 0.92 | 0.79 | 1.09 | 0.99 | 1.45 | 0 |
| | 1.2 | 0.95 | 0.95 | 1.02 | 0.78 | 0.77 | 0.89 | 1.18 | 0.81 | 0 |
| $b_{vh}$ | 0.8 | 1.07 | 1.08 | 1.09 | 0.98 | 0.8 | 0.85 | 0.86 | 0.16 | 0 |
| | 1.2 | 0.93 | 0.94 | 0.92 | 1.22 | 0.83 | 1.18 | 1.39 | 2.5 | 0 |

| | | | | | | | | | | |
|---|---|---|---|---|---|---|---|---|---|---|
| $b_{vi}$ | 0.8 | 1.06 | 1.11 | 1.06 | 0.78 | 0.92 | 1.12 | 0.96 | 0.24 | 0 |
| | 1.2 | 0.89 | 0.9 | 0.92 | 0.93 | 0.91 | 0.86 | 1.32 | 1.53 | 0 |
| $b_{iv}$ | 0.8 | 0.84 | 0.92 | 0.84 | 1.21 | 2.74 | 1.36 | 1.62 | 4.84 | 1 |
| | 1.2 | 1.09 | 1.19 | 1.06 | 0.96 | 0.77 | 1.01 | 0.84 | 0.24 | 0 |
| $d_h$ | 0.8 | -- | -- | -- | -- | -- | -- | -- | -- | 10 |
| | 1.2 | -- | -- | -- | -- | -- | -- | -- | -- | 10 |
| $d_k$ | 0.8 | 1.33 | 0.94 | 1.43 | 1.94 | 0.69 | 0.76 | 0.79 | 2.66 | 0 |
| | 1.2 | 0.64 | 1.08 | 0.54 | 1.15 | 3.66 | 2.37 | 2.06 | 5.4 | 0 |
| $d_v$ | 0.8 | 0.82 | 0.82 | 0.62 | 1.51 | 3.58 | 2.03 | 1.38 | 4.64 | 2 |
| | 1.2 | 1.27 | 1.31 | 1.13 | 0.81 | 0.63 | 0.86 | 0.69 | 0.73 | 0 |
| $d_i$ | 0.8 | 1.13 | 1.31 | 1 | 0.68 | 0.76 | 0.91 | 0.9 | 0.32 | 0 |
| | 1.2 | 0.8 | 0.78 | 0.89 | 0.91 | 2.4 | 1.47 | 1.9 | 4.97 | 4 |
| $d_h$ and $v_h$ | 0.8 | 0.96 | 1 | 0.98 | 1.05 | 1.05 | 1.07 | 1.18 | 0.89 | 0 |
| | 1.2 | 0.96 | 0.99 | 0.99 | 0.91 | 0.94 | 0.91 | 1.3 | 1.05 | 0 |
| $d_k$ and $v_k$ | 0.8 | 0.96 | 0.96 | 0.96 | 0.91 | 0.93 | 0.87 | 1.17 | 0.81 | 0 |
| | 1.2 | 1.01 | 0.99 | 1.03 | 0.84 | 0.92 | 0.92 | 1.15 | 0.56 | 0 |
| $d_i$ and $v_i$ | 0.8 | 0.93 | 0.97 | 0.81 | 1.35 | 2.53 | 1.58 | 1.47 | 4.66 | 1 |
| | 1.2 | 1.09 | 1.02 | 1.17 | 0.89 | 0.86 | 0.64 | 0.85 | 0.56 | 0 |
| $d_v$ and $v_v$ | 0.8 | 0.96 | 1.01 | 0.99 | 0.82 | 0.83 | 1.02 | 1.07 | 1.05 | 0 |
| | 1.2 | 0.96 | 0.99 | 1 | 0.87 | 0.74 | 0.93 | 1.21 | 0.81 | 0 |

**Table S4. Single parameter sensitivity analysis for parameters in morphogen and gene regulation equations.** Each parameter was perturbed up and down by 20% individually. In each case, 10 simulations were performed to calculate the resulting rhombomere lengths and SIs. Numbers of failed simulations missing at least one boundary or having >8 DCs are shown in the last column.

#### S7.2.3 Noise magnitude

Additive noise was included for both morphogen and gene regulation equations. Here we explored the effects of noise by perturbing the levels 2-fold up and down individually (**Table S5**). Rhombomere lengths were relatively stable to changes in noise level. The exception was noise in FGF signaling where larger $\mu_{f2}$ caused expansion of r3 and r5. Our choice of noise levels was effective in disturbing boundary sharpness. Some exceptions were observed for $\mu_{f1}$, $\mu_{f2}$ and $\mu_h$ at a few boundaries.

| Parameter | Fold change | Ratio of perturbed quantities to unperturbed quantities | | | | | | | | # failed simulations |
|---|---|---|---|---|---|---|---|---|---|---|
| | | r3 | r4 | r5 | SI(r2/r3) | SI(r3/r4) | SI(r4/r5) | SI(r5/r6) | DCs | |
| $\mu_{r1}$ | 0.5 | 0.96 | 1.00 | 1.01 | 0.85 | 0.74 | 0.92 | 1.20 | 1.21 | 0 |
| | 2 | 1.00 | 1.00 | 1.01 | 1.06 | 0.95 | 0.88 | 1.15 | 1.45 | 0 |
| $\mu_{r2}$ | 0.5 | 0.97 | 0.99 | 1.06 | 1.05 | 0.82 | 0.87 | 1.19 | 0.56 | 0 |
| | 2 | 1.04 | 1.02 | 1.03 | 1.12 | 1.94 | 0.99 | 1.36 | 2.50 | 0 |
| $\mu_{f1}$ | 0.5 | 0.96 | 1.00 | 0.98 | 0.87 | 0.79 | 0.81 | 1.10 | 0.73 | 0 |
| | 2 | 0.97 | 0.99 | 1.00 | 0.91 | 0.75 | 0.83 | 1.19 | 0.81 | 0 |
| $\mu_{f2}$ | 0.5 | 0.84 | 1.01 | 0.93 | 0.69 | 0.88 | 0.95 | 0.96 | 0.81 | 0 |
| | 2 | 1.44 | 0.94 | 1.35 | 4.26 | 0.75 | 0.97 | 1.83 | 5.38 | 1 |
| $\mu_v$ | 0.5 | 0.97 | 0.99 | 1.03 | 0.84 | 0.73 | 0.85 | 1.10 | 0.97 | 0 |
| | 2 | 0.96 | 1.00 | 0.99 | 0.91 | 0.78 | 0.95 | 1.09 | 0.97 | 0 |
| $\mu_v$ | 0.5 | 0.98 | 1.03 | 1.03 | 0.99 | 0.61 | 0.71 | 0.62 | 0.73 | 0 |
| | 2 | 0.95 | 0.88 | 1.11 | 1.03 | 1.15 | 1.91 | 1.52 | 2.15 | 1 |
| $\mu_h$ | 0.5 | 0.95 | 1.00 | 0.96 | 1.16 | 0.69 | 0.96 | 1.18 | 0.73 | 0 |

| | | | | | | | | | | |
|---|---|---|---|---|---|---|---|---|---|---|
| | 2 | 1.02 | 1.01 | 1.06 | 1.01 | 0.82 | 0.76 | 0.86 | 0.97 | 0 |
| $\mu_k$ | 0.5 | 1.00 | 0.99 | 1.02 | 1.04 | 1.00 | 0.94 | 1.09 | 0.97 | 0 |
| | 2 | 0.92 | 0.97 | 0.93 | 1.22 | 1.83 | 1.22 | 1.47 | 3.39 | 0 |

**Table S5. Noise magnitude and rhombomere formation.** Noise levels were perturbed 2-fold up and down and 10 simulations were performed to calculate each rhombomere length and SI. Numbers of failed simulations missing at least one boundary or having >8 DCs are shown in the last column.

**S7.2.4 Cell mechanical strength and its coupling with gene regulation**

The cell mechanical model relies on many parameters in **Table S1**. Their values are based on previous work [1]. We assess sensitivity with respect to the strength of the critical features in the mechanical model, such as the ratio between attraction and repulsion and the strength/timescale of the cell sorting.

We investigate how the model performs with different attraction-repulsion ratios by either perturbing attraction or repulsion strength individually (**Table S6**). Specifically, in Eq. (S7), we perturb attraction by perturbing $U_{\text{inter}}^{\text{Atr}}$ and $V_{\text{inter}}^{\text{Atr}}$ equally or repulsion by perturbing $U_{\text{inter}}^{\text{Rep}}$ alone. We measured the effects of perturbations on lengths of three rhombomeres, SIs of four boundaries and number of DCs. We calculate the ratio between quantities in perturbed cases and quantities in unperturbed cases at the end of the simulation (14 hpf). Rhombomere lengths are relatively insensitive to changes in mechanical forces since cell fates are mainly regulated by morphogens and gene expression. The SIs are positively related to the attraction-repulsion ratio, where a smaller ratio leads to sharper boundaries. However, the valid range of the attraction-repulsion ratio to generate clear five-segment patterns is narrow (~10%, an example in **Fig S7A**), where a 20% reduction on attraction-repulsion ratio leads to separation and gaps of cells with different identities (**Table S6** and **Fig S7B**) and a 20% increase of attraction-repulsion ratio leads to rougher boundaries (**Fig S7C**).

We also examined how the coupling between cell mechanics and gene regulation affects the patterns. We perturbed the strength/timescale of the cell mechanical model. The rhombomere lengths are insensitive to this perturbation since gene regulation plays a major role in controlling rhombomere lengths. Because cell sorting plays major roles in boundary refinement, increased mechanical force strength makes boundaries sharper with smaller SIs and reduced numbers of DCs. This result supports the conclusion that the combination of global (gene regulation) and local (cell sorting) processes works synergistically in boundary sharpening.

| **Components** | **Fold change** | **Ratio of perturbed quantities to unperturbed quantities** | | | | | | | | **# failed simulations** |
|---|---|---|---|---|---|---|---|---|---|---|
| | | **r3** | **r4** | **r5** | **SI(r2/r3)** | **SI(r3/r4)** | **SI(r4/r5)** | **SI(r5/r6)** | **DCs** | |
| Attractions | 0.8* | 1.31 | 1.00 | 1.15 | 0.00 | 0.22 | 0.20 | 0.18 | 0.65 | 0 |
| | 0.9 | 0.97 | 1.02 | 1.01 | 0.41 | 0.61 | 0.47 | 0.96 | 0.65 | 0 |
| | 1.1 | 0.94 | 1.00 | 0.98 | 1.23 | 1.28 | 1.61 | 1.30 | 1.45 | 0 |
| | 1.2 | 0.95 | 0.97 | 0.99 | 1.93 | 2.29 | 2.53 | 2.06 | 2.18 | 0 |
| Repulsions | 0.8 | 1.01 | 0.92 | 1.01 | 2.59 | 3.05 | 3.61 | 2.68 | 2.66 | 0 |
| | 0.9 | 0.96 | 0.97 | 1.01 | 1.44 | 1.25 | 1.86 | 1.50 | 1.53 | 0 |
| | 1.1 | 0.95 | 1.02 | 1.01 | 0.48 | 0.55 | 0.56 | 0.89 | 0.56 | 0 |
| | 1.2* | 1.13 | 1.00 | 1.16 | 0.15 | 0.28 | 0.36 | 0.17 | 0.73 | 0 |
| Cell mechanics strength | 0.8 | 0.97 | 1.01 | 0.98 | 1.02 | 0.83 | 1.04 | 1.16 | 1.05 | 0 |
| | 0.9 | 0.95 | 0.99 | 0.98 | 0.85 | 0.84 | 0.85 | 1.08 | 0.89 | 0 |
| | 1.1 | 0.95 | 1.02 | 0.98 | 0.81 | 0.72 | 0.99 | 1.12 | 0.65 | 0 |
| | 1.2 | 0.95 | 0.99 | 1.02 | 0.71 | 0.67 | 0.95 | 1.17 | 1.13 | 0 |

**Table S6**. **Attractive and repulsive forces in rhombomere formation**. Attraction or repulsion was perturbed up or down by 10% and 20%, either individually or together. In each case, 10 simulations were performed to calculate the statistics for rhombomere lengths and SIs. Numbers of failed simulations missing at least one boundary or having >8 DCs are shown in the last column. Note: *Simulations show big gaps between clones of cells with different identities.

### S8. Experimental measurements of hindbrain dimensions

Here we list the measurements of hindbrain dimensions made using whole mount in situ hybridization at 11-14 hpf (**Fig 1C**). At 11 hpf, the *krox20* expression domain in r3 has rough, intermittent boundaries. Thus, at this stage we only measured total hindbrain length and the RA production region. In some samples it was difficult to define the RA production region, and these were omitted in calculating average lengths.

| Time | Sample NO. | LR width ($L_2(t)$; µm) | AP lengths (µm) | | | | | | |
|---|---|---|---|---|---|---|---|---|---|
| | | | MHB to r3 | r3 | r3-r5 | r5 | r5 to RA production region | Hindbrain Total Length ($p(t)$) | RA production region length |
| 11hpf | 1 | 255 | -- | -- | -- | -- | -- | 343 | 278 |
| | 2 | 237 | -- | -- | -- | -- | -- | 338 | 273 |
| | 3 | 308 | -- | -- | -- | -- | -- | 281 | 284 |
| | 4 | 312 | -- | -- | -- | -- | -- | 294 | -- |
| | 5 | 269 | -- | -- | -- | -- | -- | 313 | 302 |

| | | | | | | | | | |
|---|---|---|---|---|---|---|---|---|---|
| | 6 | 303 | -- | -- | -- | -- | -- | 355 | 306 |
| | 7 | 316 | -- | -- | -- | -- | -- | 343 | 281 |
| | 8 | 272 | -- | -- | -- | -- | -- | 339 | -- |
| | 9 | 278 | -- | -- | -- | -- | -- | 298 | 281 |
| | **Mean** | **283** | **--** | **--** | **--** | **--** | **--** | **323** | **286** |
| 12hpf | 1 | 206 | 74 | 32 | 33 | 27 | 273 | 440 | -- |
| | 2 | 152 | 64 | 34 | 33 | 40 | 284 | 455 | 206 |
| | 3 | 139 | 71 | 43 | 28 | 49 | 298 | 490 | 203 |
| | 4 | 182 | 74 | 38 | 35 | 31 | 283 | 460 | 209 |
| | 5 | 175 | 59 | 31 | 30 | 37 | 252 | 410 | 179 |
| | 6 | 154 | 61 | 28 | 28 | 37 | 260 | 414 | 217 |
| | 7 | 152 | 68 | 36 | 30 | 31 | 262 | 427 | 213 |
| | 8 | 150 | 67 | 39 | 32 | 42 | 277 | 457 | 206 |
| | 9 | 153 | 75 | 35 | 34 | 39 | 274 | 457 | 200 |
| | **Mean** | **162** | **68** | **35** | **31** | **37** | **274** | **445** | **233** |
| 13hpf | 1 | 109 | 53 | 33 | 31 | 35 | 219 | 370 | 245 |
| | 2 | 131 | 72 | 40 | 31 | 44 | 236 | 423 | 357 |
| | 3 | 186 | 91 | 47 | 30 | 54 | 274 | 497 | 369 |
| | 4 | 162 | 81 | 41 | 37 | 35 | 266 | 459 | 308 |
| | 5 | 133 | 63 | 29 | 20 | 34 | 276 | 422 | 260 |
| | 6 | 123 | 86 | 43 | 42 | 41 | 246 | 458 | 348 |
| | 7 | 127 | 67 | 59 | 29 | 49 | 242 | 446 | 354 |
| | 8 | 157 | 88 | 42 | 31 | 34 | 256 | 452 | 271 |
| | 9 | 128 | 67 | 41 | 33 | 39 | 257 | 437 | 280 |
| | **Mean** | **140** | **74** | **42** | **32** | **40** | **252** | **440** | **310** |
| 14hpf | 1 | 109 | 85 | 39 | 38 | 36 | 230 | 428 | -- |
| | 2 | 110 | 80 | 48 | 29 | 33 | 241 | 431 | 365 |

| | | | | | | | | |
|---|---|---|---|---|---|---|---|---|
| | 3 | 96 | 82 | 36 | 45 | 41 | 198 | 402 | 420 |
| | 4 | 112 | 71 | 46 | 29 | 34 | 234 | 414 | 375 |
| | 5 | 97 | 93 | 43 | 35 | 43 | 236 | 450 | 452 |
| | 6 | 99 | 74 | 46 | 33 | 36 | 233 | 423 | -- |
| | 7 | 106 | 62 | 46 | 31 | 38 | 214 | 391 | 375 |
| | 8 | 111 | 92 | 41 | 31 | 38 | 223 | 426 | 379 |
| | 9 | 91 | 65 | 32 | 29 | 28 | 190 | 343 | -- |
| | **Mean** | **104** | **78** | **42** | **33** | **37** | **222** | **412** | **394** |

**Table S7. Experimental measurement for hindbrain dimensions.**

#### S9. Multiplicative noise in gene regulation models

Additive noise in morphogen (Eq. (5) and Eq. (8)) and gene regulation equations (Eq. (10) was used throughout most of this study. Here we investigate multiplicative noise. The equations for RA with multiplicative noise (bold font) are given as the following:

$$\begin{aligned}
\frac{\partial [RA]_{out}}{\partial t} + \nabla \cdot \left( \boldsymbol{V}[RA]_{out} \right) &= D_r \Delta [RA]_{out} + A(\boldsymbol{x}, t) \\
&\quad + k_r [RA]_{in} - (1+\beta_r) k_r [RA]_{out} \boldsymbol{+ h_{r1} [RA]_{out} \frac{dm_{r1}(t)}{dt}}, \\
\frac{\partial [RA]_{in}}{\partial t} + \nabla \cdot \left( \boldsymbol{V}[RA]_{in} \right) &= k_r [RA]_{out} - k_r [RA]_{in} - d_r (x^{(1)}) [RA]_{in} \\
&\quad \boldsymbol{+ h_{r2} [RA]_{in} \frac{dm_{r2}(t)}{dt}},
\end{aligned} \tag{S20}$$

where $h_{r1}[RA]_{out} \frac{dm_{r1}(t)}{dt}$ and $h_{r2}[RA]_{in} \frac{dm_{r2}(t)}{dt}$ are the multiplicative white noise. Similarly, equations for FGF and morphogen downstream genes with multiplicative noise are given:

$$\frac{\partial[Fgf]_{free}}{\partial t}+\nabla\cdot\left(\boldsymbol{V}[Fgf]_{free}\right)=D_f\Delta[Fgf]_{free}+A_f([H],\boldsymbol{x})+k_{rf}[Fgf]_{signal}$$
$$-(d_{f1}+k_f)[Fgf]_{free}\boldsymbol{+h_{f1}[Fgf]_{free}}\frac{\boldsymbol{dm_{f1}(t)}}{\boldsymbol{dt}},$$
$$\frac{\partial[Fgf]_{signal}}{\partial t}+\nabla\cdot\left(\boldsymbol{V}[Fgf]_{signal}\right)=k_f[Fgf]_{free}-k_{rf}[Fgf]_{signal}$$
$$-d_{f2}[Fgf]_{signal}\boldsymbol{+h_{f2}[Fgf]_{signal}}\frac{\boldsymbol{dm_{f2}(t)}}{\boldsymbol{dt}}, \tag{S21}$$

$$\frac{d[H]_i}{dt}=v_h\frac{a_{hh}[H]_i^2+a_{rh}[RA]_{in}^2}{1+a_{hh}[H]_i^2+a_{rh}[RA]_{in}^2+b_{kh}[K]_i^2+b_{vh}[V]_i^2}-d_h[H]_i\boldsymbol{+h_h[H]_i}\frac{\boldsymbol{dm_h}}{\boldsymbol{dt}},$$
$$\frac{d[K]_i}{dt}=v_k\frac{a_{kk}[K]_i^2+a_{fk}[Fgf]_{signal}^2}{1+a_{kk}[K]_i^2+a_{fk}[Fgf]_{signal}^2+b_{hk}[H]_i^2}-d_k[K]_i\boldsymbol{+h_k[K]_i}\frac{\boldsymbol{dm_k}}{\boldsymbol{dt}},$$
$$\frac{d[V]_i}{dt}=v_v\frac{a_{rv}[RA]_{in}^2}{1+a_{rv}[RA]_{in}^2+b_{iv}[I]_i^2}-d_v[V]_i\boldsymbol{+h_v[V]_i}\frac{\boldsymbol{dm_v}}{\boldsymbol{dt}},$$
$$\frac{d[I]_i}{dt}=v_i\frac{1}{1+b_{vi}[V]_i^2}-d_i[I]_i\boldsymbol{+h_i[I]_i}\frac{\boldsymbol{dm_i}}{\boldsymbol{dt}}. \tag{S22}$$

Compared with additive noise (**Fig 4)**, our findings suggest that the positive effects of rapid initial convergence on segmental pattern robustness also occur with multiplicative noise (**Fig S8**). Initial rapid convergence results in the smallest r4 A-P length compared with medium or slow initial convergence. The initial slow convergence has largest rhombomere lengths for r3, r4 and r5. All four boundaries are also sharper with initial rapid convergence compared with slow initial convergence. In addition, rapid initial convergence results in fewer numbers of DCs compared to slow initial convergence. Medium convergence leads to sharper boundaries for r3/r4, r4/r5 and r5/r6 than rapid initial convergence, but leads to defects in rhombomere lengths, with r4 larger than r3 and r5, and r5 is smaller than r3. A difference between the effects of multiplicative versus additive noise is that with the former, the posterior boundaries (i.e. r4/r5 and r5/r6) appear much rougher than anterior boundaries (r2/r3 and r3/r4). This is likely because RA levels are higher posteriorly and RA noise levels are also higher with multiplicative noise. Noise magnitudes used in **Fig S8** are provided in **Table S8**.

| Multiplicative noise magnitude | values |
|---|---|
| $h_{r1}$ | 0.002 |
| $h_{r2}$ | 0.002 |
| $h_{f1}$ | 0.002 |
| $h_{f2}$ | 0.002 |
| $h_v$ | 0.02 |
| $h_i$ | 0.02 |
| $h_k$ | 0.02 |
| $h_h$ | 0.02 |

**Table S8. Multiplicative noise magnitudes used in Fig S8.**

**S10. Model with cell proliferation and growth**

In the patterning period we have modeled, cells divide in the hindbrain. Cell proliferation and growth are coupled with cell migration along the D-V axis that thickens the neural plane. As a result, in the two-dimensional plane consisting of A-P and L-R axes, the number of cells remains relatively unchanged [5]. We therefore simplified our two-dimensional model by not considering cell proliferation and growth.

Between 11-14 hpf in zebrafish embryos, cells begin to enter the $16^{th}$ cell cycle and divide asynchronously with an average cell cycle length of 4 hours [9]. Cells tend to divide and separate in the direction perpendicular to the A-P axis [9]. We modeled cell division according to the experimental evidence on cell cycle and division orientation. In the model, we randomly assigned division times for each cell in a 4-hour window (11-15 hpf), where a cell would not divide if it had not divided by 14 hpf (**Fig S9C**). For each cell, division has three key stages: elongation, separation and growth (**Fig S9A**). First, a cell needs to determine its division line. We modeled each cell with a hexagonal shape. The division line goes through mid-points of two opposite sides of each hexagon. With three pairs of opposite sides, there are three possible choices of division line. We selected the division line having the minimum angle relative to the A-P axis (**Fig S9A**). After the division line is selected, the intrinsic lengths in Eq. (S11) for the sides intersected by the line of orientation grow linearly with similar velocities. Cell elongation will continue for 30 minutes until the lengths of the growing sides increase $l_{out}$. Then the cell divides into two daughter cells and the intrinsic length $l_{ij}$ in Eq. (S11) for all sides becomes 60% of their default values. The two daughter cells grow to normal size in 30 minutes with linearly growing intrinsic lengths (**Fig S9A**).

Similar to **Fig 4**, we perform simulations to compare the effects of convergence rates on pattern formation (**Fig S9B** and **S9D-F**). Our major finding, that rapid initial convergence improves pattern robustness, remains. A clear five-segment pattern is established (**Fig S9B**), but cells distribute more tightly than in cases without cell division. The lengths of all three rhombomeres under three different convergence patterns (**Fig S9D**) are similar to the cases without division (**Fig 4G**), but have higher variance. Rapid initial convergence yields the smallest SIs for all four boundaries with both medium and slow initial convergence rates (**Fig S9E**).

**S11. Models without convergent extension**

Without convergent extension, morphogens change less because the tissue no longer elongates to scale morphogens distributions **(Fig S10)**. Indeed, most cells commit to fates much earlier and the five-segment pattern is established around 12 hpf (**Fig S10A**). From 12-14 hpf, boundaries are refined through cell sorting, though some minor refinements of boundaries are observed. Comparisons between models with and without convergent extension (**Fig S10B-D**) reveal that without convergent extension, much smaller r3 and r5 A-P lengths result and all boundaries are rougher with higher SIs. r3 and r5 are only one or two cells long along the A-P axis, and such a limited number of neighboring cells with similar identities provides very weak adhesion and difficulty sorting cells back to their correct segments (**Fig S10A).** These observations further support the idea of a trade-off between segment size and boundary sharpness.

**S12. Models without advections of intracellular morphogens**

As the tissue deforms, the extra-cellular morphogens may have active motion driven by the tissue dynamics while the intra-cellular signals induced by morphogens may also move due to the cell movements in Eulerian coordinates. In a continuum model using Eulerian coordinates, the advections are usually required to capture the morphogen dynamics with moving boundaries [10].

We also provide a model without advection for the intracellular morphogens for both RA and FGF, using the modified equations of Eq. (5) and (9):

$$\begin{aligned}\frac{\partial[RA]_{in}}{\partial t} &= k_r[RA]_{out} - k_r[RA]_{in} - d_r(x^{(1)})[RA]_{in} + \mu_{r2}\frac{dw_{r2}(t)}{dt},\\ \frac{\partial[Fgf]_{signal}}{\partial t} &= k_f[Fgf]_{free} - k_{rf}[Fgf]_{signal} - d_{f2}[Fgf]_{signal} + \mu_{f2}\frac{dw_{f2}(t)}{dt}.\end{aligned} \tag{S23}$$

Our main result showing positive effects of rapid initial convergence on patterning remains when such advection is removed. The rapid initial convergence induces sharper boundaries over all four boundaries and fewer DCs than medium and slow initial convergence (**Fig S11C** and **S11D**). The discrepancy in intracellular RA between rapid and slow initial convergence decreases when advection is removed but still exists (**Fig S11E** and **S11F**).
