## Supplementary Figures for "Multiple morphogens and rapid elongation promote segmental patterning during development"

### Supplementary Figures S1-S10

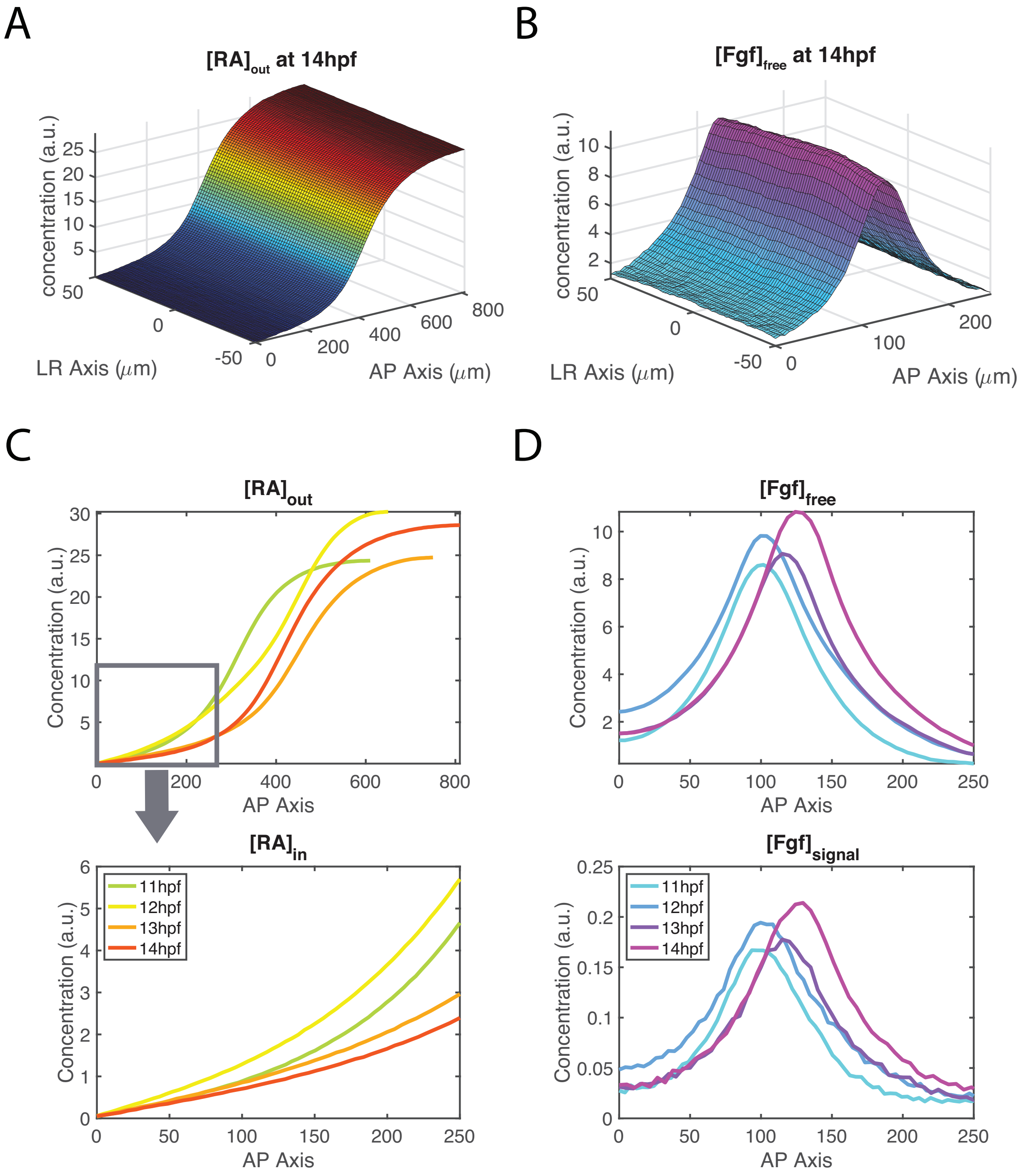


**Fig S1 Morphogen distributions. (A)** The distribution of extracellular RA $[RA]_{out}$ at 14 hpf. **(B)** The distribution of free diffusible FGF, $[Fgf]_{free}$ at 14 hpf. **(C)** The A-P distribution of $[RA]_{out}$ in the morphogen domain and the higher magnification A-P distribution of $[RA]_{in}$ for the A-P range present in the responding tissue domain. The curves show the average morphogen level along the L-R axis. **(D)** The A-P distribution of free diffusible FGF, $[Fgf]_{signal}$, FGF signal, $[Fgf]_{signal}$. The curves show the average morphogen levels along the L-R axis.

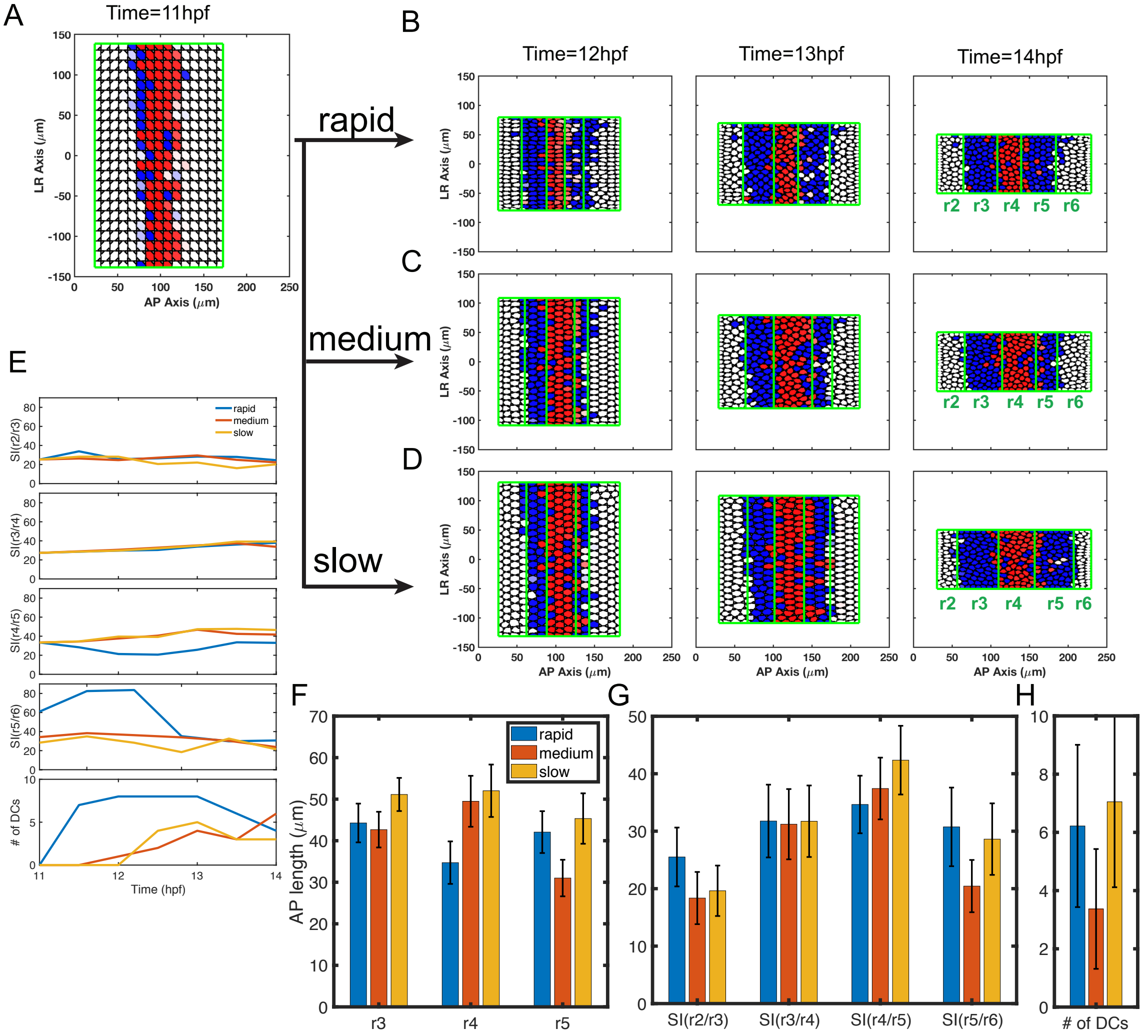


**Fig S2. Simulations with gene regulation alone with different convergence rates.** **(A-D)** Time series of cell distributions with different convergence rates from 11-14 hpf. *hoxb1a* (red) and *krox20* (blue) expression. **(A)** Three simulations start with the same initial cell distribution (11 hpf) generated by the gene expression model. Cell distributions with **(B)** rapid, **(C)** medium and **(D)** slow initial convergence rates from 12-14 hpf. **(E)** The boundary sharpness index (SI) for four boundaries (SI(r2/r3), SI(r3/r4), SI(r4/r5) and SI(r5/r6)) and number of DC versus time. Histograms depicting three convergence rates analyzed for **(F)** rhombomere lengths of r3-5, **(G)** SI and **(H)** the number of DC. 100 independent stochastic simulations for each convergence rate based on the same parameters. Error bars represent standard deviation.

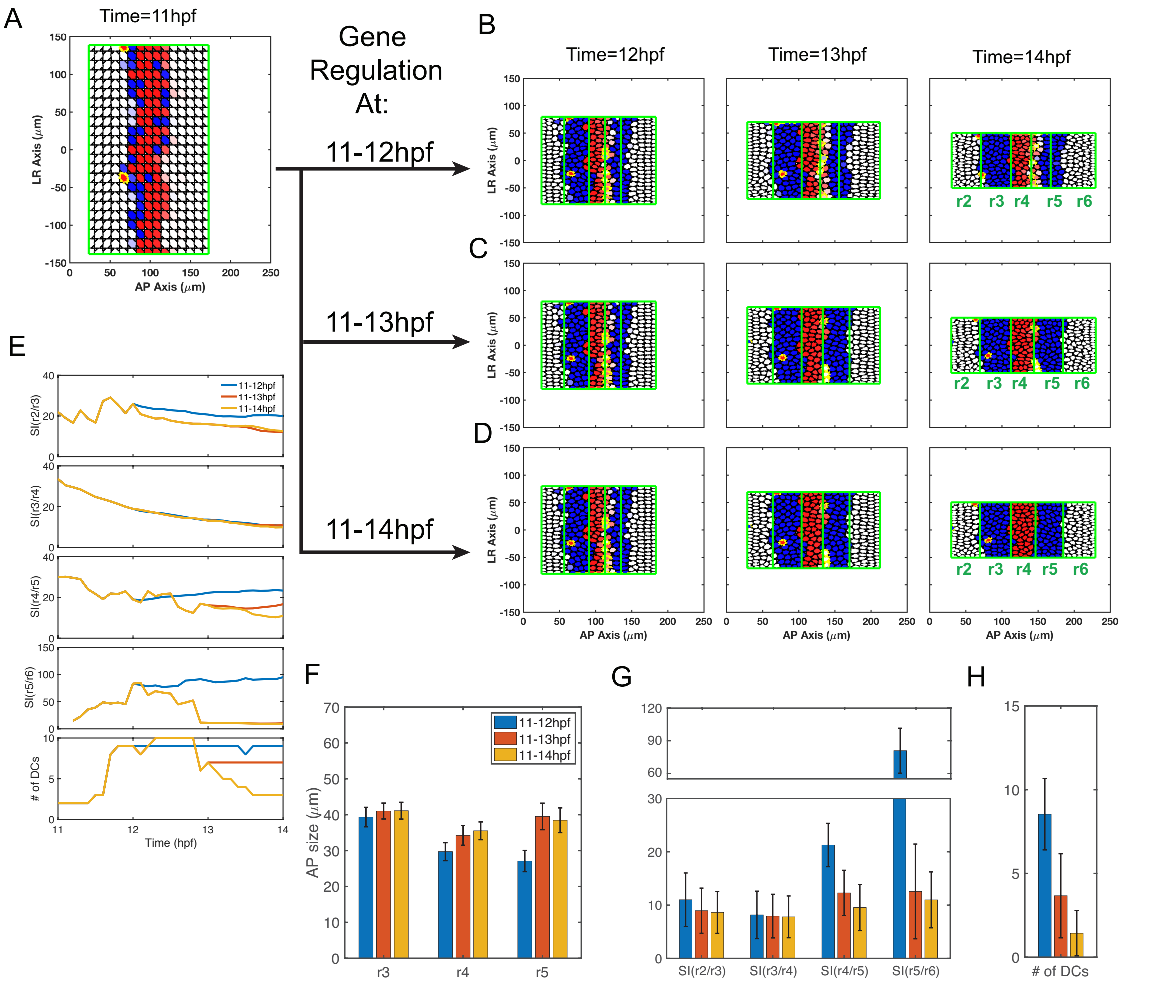


**Fig S3. Timing of gene regulatory interactions influences hindbrain patterning. (A)** Initial cell distribution at 11 hpf. **(B-D)** Time series of cell distributions with gene regulation limited to 1, 2 or 3 hour periods from 11-14 hpf: **(B)** 11-12 hpf, **(C)** 11-13 hpf and **(D)** 11-14 hpf. All start with the same initial cell distribution and gene expression pattern. *hoxb1a* expression (red); *krox20* expression (blue). Dislocated cells (DCs) are highlighted by yellow edges. **(E)** The sharpness index (SI) for four boundaries (SI(r2/r3), SI(r3/r4), SI(r4/r5) and SI(r5/r6)) and number of DCs versus time. **(F)** Rhombomere lengths, **(G)** SIs and **(H)** DCs. 100 independent stochastic simulations for each type of convergence were computed under the same parameters. Error bars represent the standard deviation.

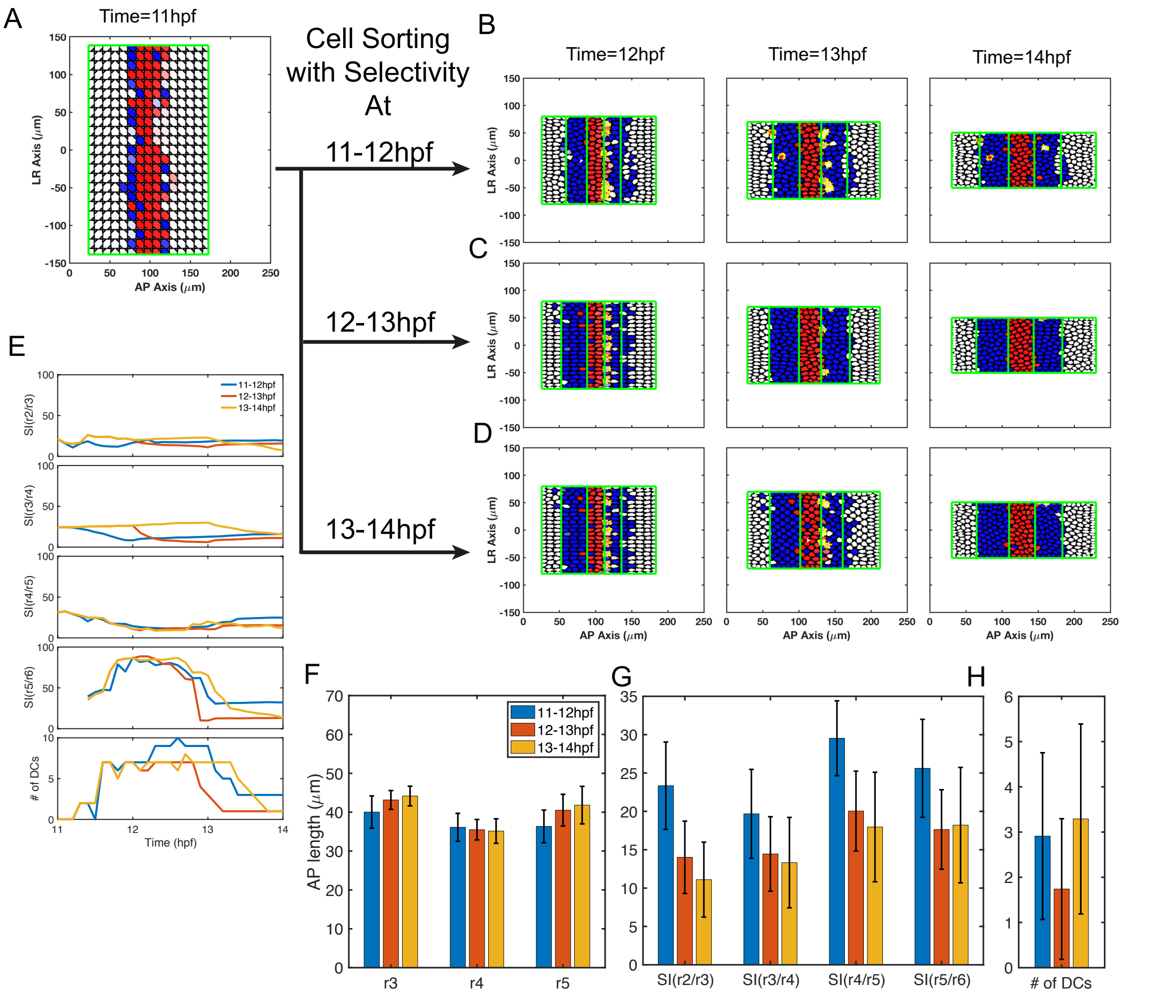


**Fig S4. Timing of selective cell sorting influences hindbrain patterning. (A)** Initial cell distributions at 11 hpf. **(B-D)** Time series of cell distributions with selective cell sorting during three different one-hour periods: **(B)** 11-12 hpf, **(C)** 12-13 hpf and **(D)** 13-14 hpf. All three simulations start with the same initial cell distribution and gene expression pattern (11 hpf) generated by gene expression models. Selectivity is removed by setting the similarity function as a constant $\mathcal{F} = 0.5$ such that cells sort randomly. *hoxb1a* expression (red), *krox20* expression (blue). Dislocated cells (DCs) are highlighted by yellow edges. **(E)** Sharpness indices (SIs) and numbers of DCs (versus time. **(F)** Rhombomere lengths, **(G)** SIs and **(H)** DCs. 100 independent stochastic simulations for each type of convergence were computed under the same parameters. Error bars represent the standard deviation.

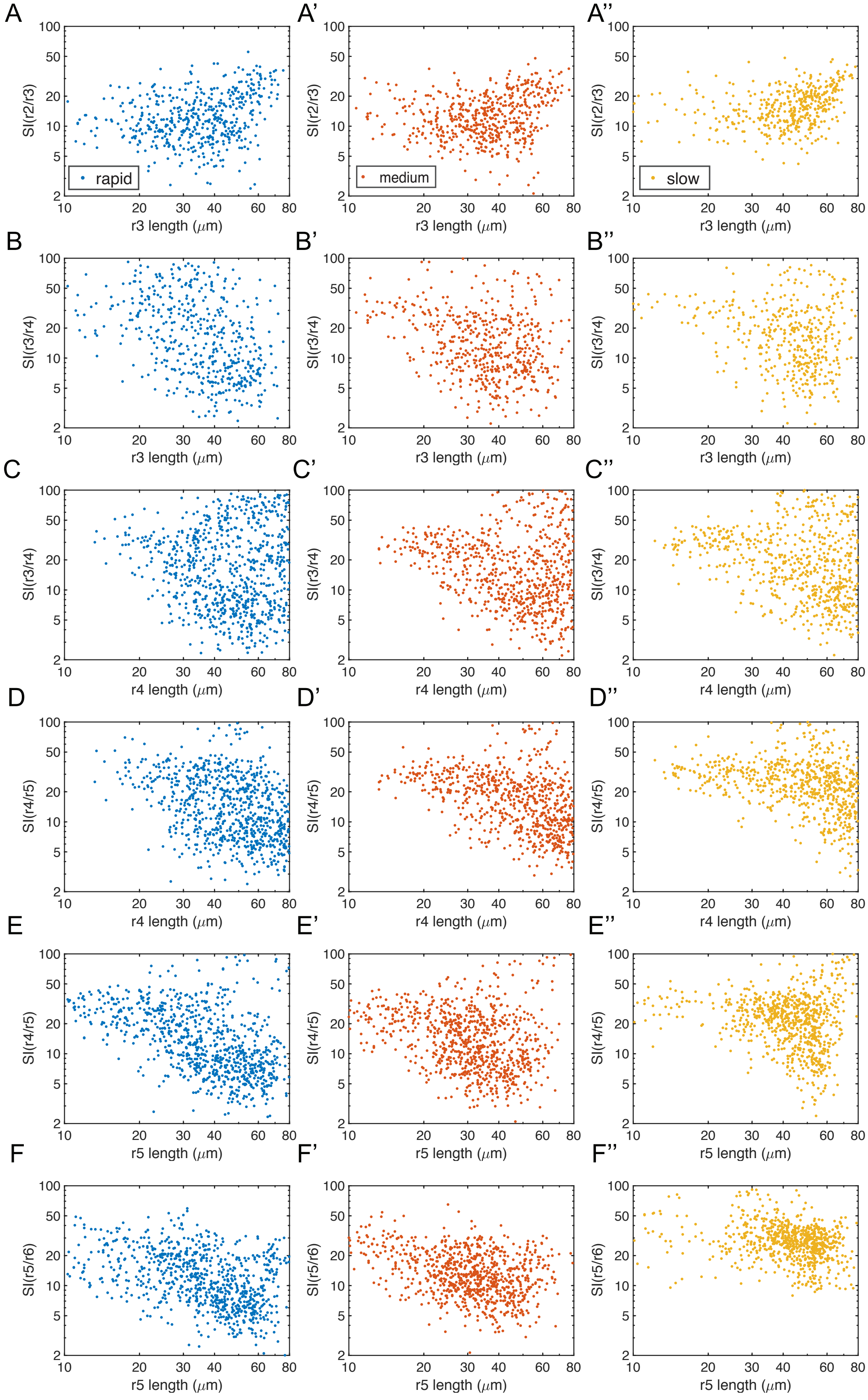
A
A'
A''
rapid
medium
slow
SI(r2/r3)
r3 length (μm)
B
B'
B''
SI(r3/r4)
r3 length (μm)
C
C'
C''
SI(r3/r4)
r4 length (μm)
D
D'
D''
SI(r4/r5)
r4 length (μm)
E
E'
E''
SI(r4/r5)
r5 length (μm)
F
F'
F''
SI(r5/r6)
r5 length (μm)
100
50
20
10
5
2
10
20
30
40
60
80

**Fig S5. The relations between rhombomere length and the level of boundary sharpness (SI).** Parameters for gene regulation are randomly perturbed and a total of n=1000 simulations are displayed for each convergence rate. There are 513, 563 and 452 simulations for rapid, medium and slow convergence, respectively, that successfully generate the r2-6 pattern with four boundaries. Each point represents the corresponding quantities for each simulation. **(A-A'')** Length of r3 versus SI(r2/r3): **(A)** rapid, **(A')** medium and **(A'')** slow convergence. **(B-B'')** Length of r3 versus SI(r3/r4): **(B)** rapid, **(B')** medium and **(B'')** slow convergence. **(C-C'')** Length of r4 versus SI(r3/r4): **(C)** rapid, **(C')** medium and **(C'')** slow convergence. **(D-D'')** Length of r4 versus SI(r4/r5): **(D)** rapid, **(D')** medium and **(D'')** slow convergence. **(E-E'')** Length of r5 versus SI(r4/r5): **(E)** rapid, **(E')** medium and **(E'')** slow convergence. **(F-F'')** Length of r5 versus SI(r5/r6): **(F)** rapid, **(F')** medium and **(F'')** slow convergence.

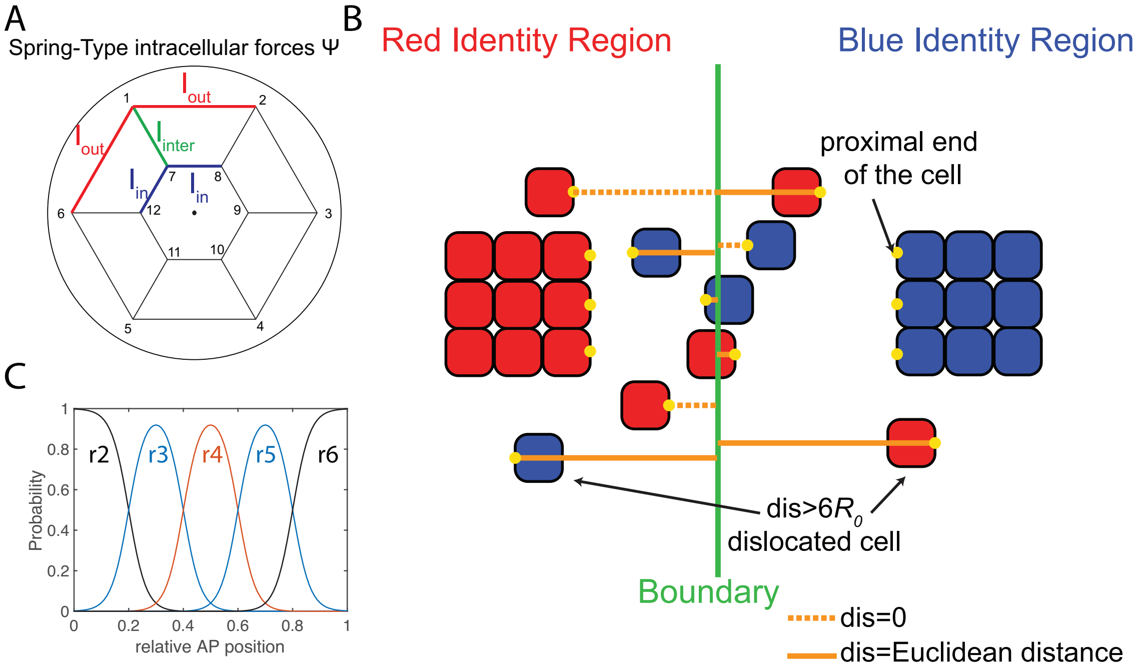


**Fig S6. Additional information for the model. (A).** Cell representation in SCEM: Each cell is represented by two layers of nodes and each layer has $N_{node}$=6 nodes with a hexagonal structure. The pairwise lengths used in Eq. (S9) have three possible constant values, $l_{out}$, $l_{in}$, $l_{inter}$. The length between two neighbor nodes in the outer layer is $l_{out}$. The length between two neighbor nodes in the inner layer is $l_{in}$. For two nodes in different layers with same angle, the length is $l_{inter}$. **(B).** An illustration of the distance function between cells and the boundary. For cells with identities belonging to the left side of the boundary (red cells), the proximal end is taken as the right-most edge of the cell. The distance is the Euclidean distance from the proximal edge to the boundary. Similarly, for the cell with identity belonging to the right side of the boundary (blue cells), the proximal edge is taken as the left-most edge of the cell. The distance is the Euclidean distance from the proximal edge to the boundary. For distances greater than 6*r, (r is the radius of the cell), the cell is identified as a dislocated cell (DC). **(C)** The probability distribution for generating the initial distribution of cells in Fig 4B. There are five cell identities and each of them is a Gaussian with respect to A-P position.

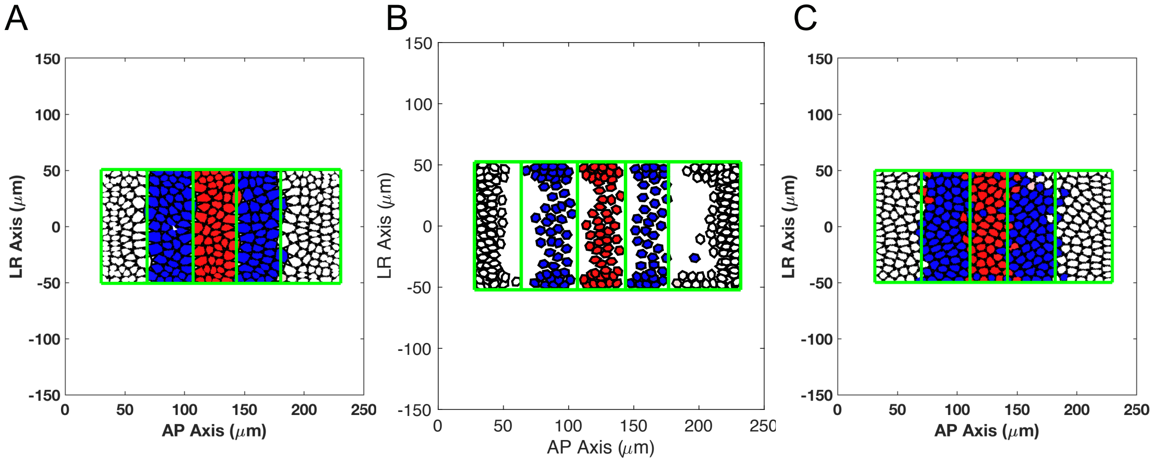


**Fig S7. Perturbations on mechanical model.** The cell distributions at the end of the simulation (14hpf) are shown for the case with perturbed mechanical forces. **(A)** 10% reduction on attraction results in five-segment pattern with sharp boundaries. **(B)** 20% reduction on attraction results in completely separation and gaps between cells with different identities. Cells with same identities gather together to form clones. The model fails to mimic the pattern formations. **(C)** 20% reduction on repulsion results in five-segment pattern with mixture boundaries.

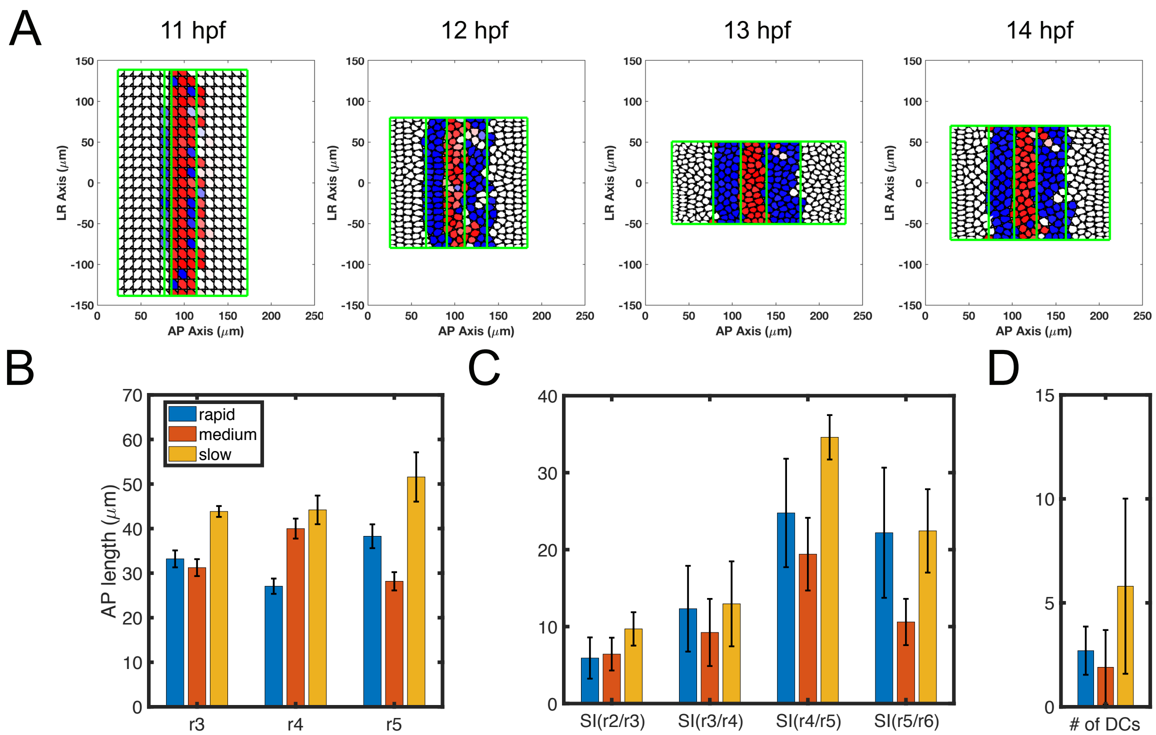


**Fig S8. Simulations with multiplicative noise**. **(A)** Time series of cell distribution with rapid initial convergence. **(B-D)** Comparisons of rapid early convergence, medium convergence and slow initial convergence. Statistics for **(B)** rhombomere lengths, **(C)** SIs and **(D)** DCs. 10 independent stochastic simulations for each type of convergence were computed under the same parameters. Error bars represent the standard deviation.

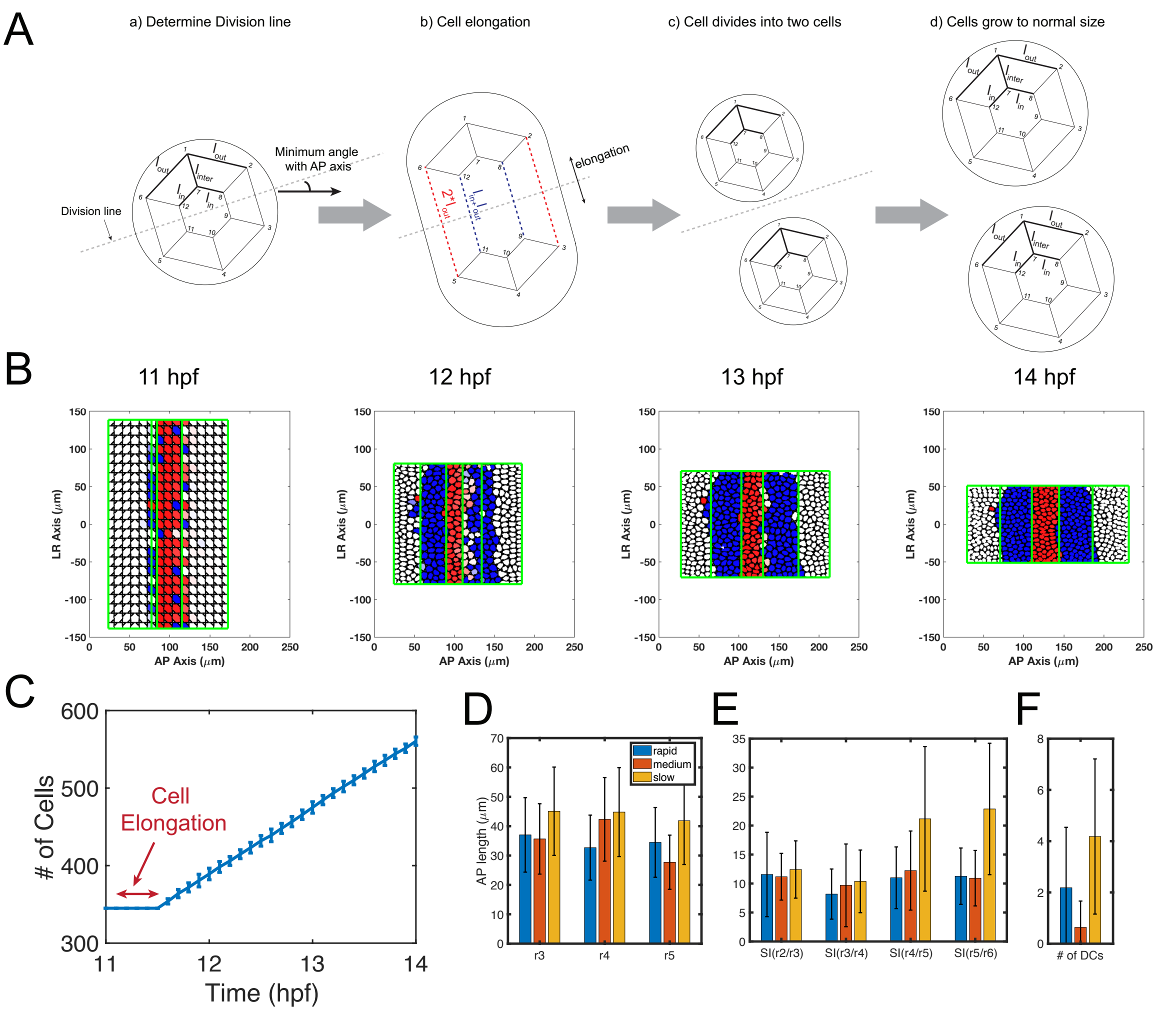


**Fig S9**. **Simulations with cell proliferation and growth.** (A) A illustration of how a cell divides. First, the division line is selected and cell elongates in the direction perpendicular to the division line. When the cell elongates to a certain length, it divides and separate into two daughter cells. Then the two daughter cells grow to normal cells size. **(B)** Time series of cell distribution with rapid initial convergence. **(C)** The number of cells over time (11-14hpf). The curve is from 10 independent simulations and error bar represent standard deviation. There are no cell divides during in 11-11.5 hpf because cells need 30 minutes to prepare for division (cell elongation) and there are no cells elongated before our initial simulation time 11hpf. **(D-F)** Comparisons of rapid early convergence, medium convergence and slow initial convergence. Statistics for **(D)** rhombomere lengths, **(E)** SIs and **(F)** DCs. 10 independent stochastic simulations for each type of convergence were computed under the same parameters. Error bars represent the standard deviation.

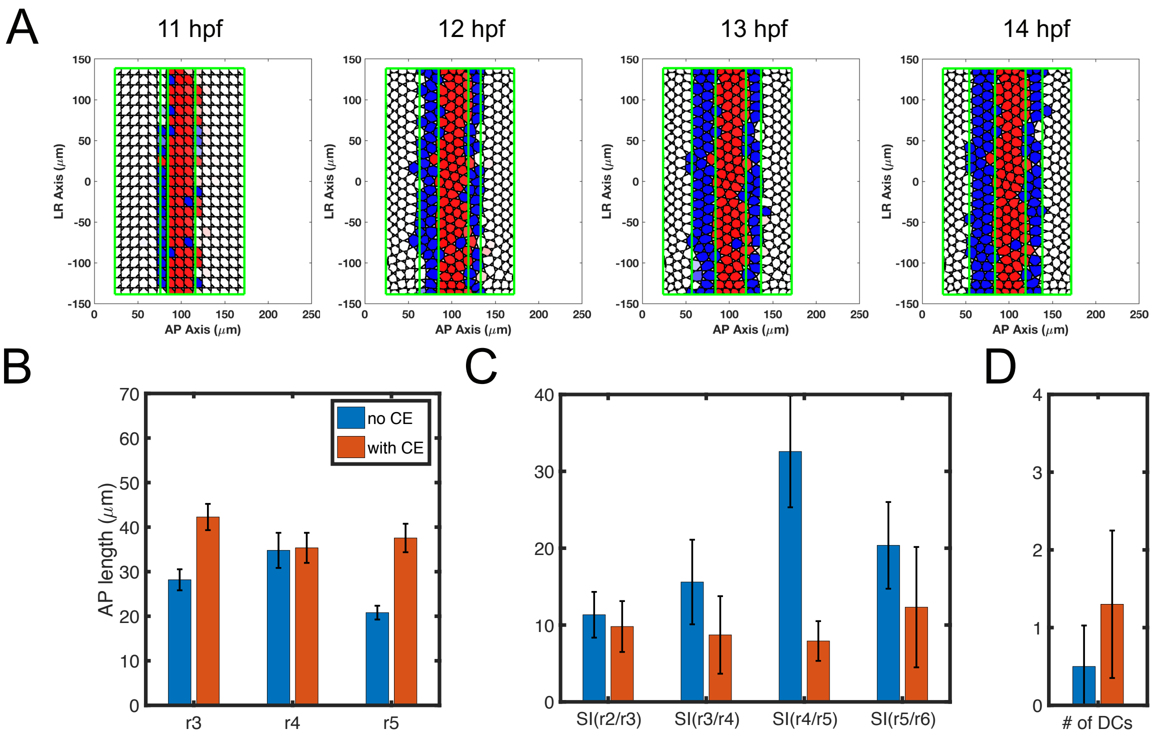


**Fig S10. Comparison for models with convergent extension (CE) and without convergent extension. (A)** Time series of cell distribution with rapid initial convergence. **(B-D)** Comparisons for models without convergent extension and with convergent extesnion. Statistics for **(B)** rhombomere lengths, **(C)** SIs and **(D)** DCs. 100 independent stochastic simulations for each type of convergence were computed under the same parameters. Error bars represent the standard deviation.

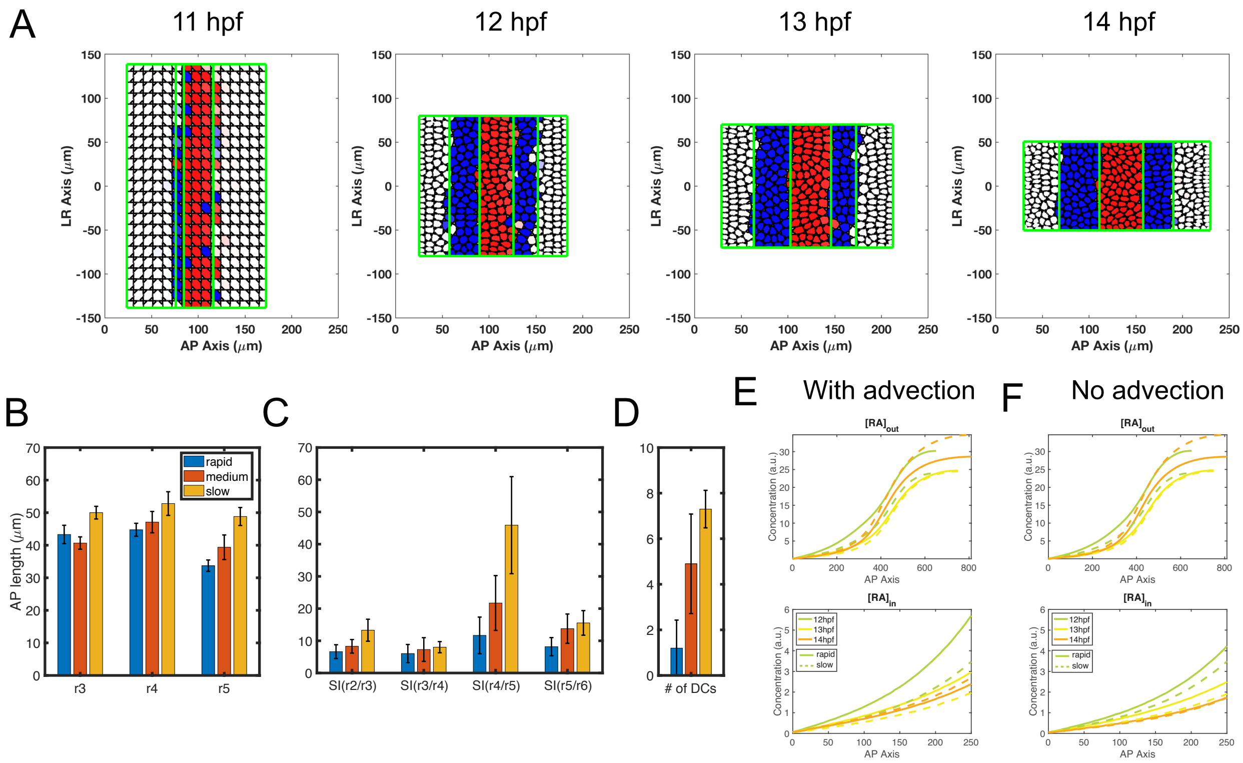


**Fig. S11. Simulations for equations without advections on $[RA]_{in}$ and $[FGF]_{signal}$. (A)** Time series of cell distribution with rapid initial convergence. **(B-D)** Comparisons of rapid early convergence, medium convergence and slow initial convergence. Statistics for **(B)** rhombomere lengths, **(C)** SIs and **(D)** DCs. 10 independent stochastic simulations for each type of convergence were computed under the same parameters. Error bars represent the standard deviation. **(E-F)** the A-P distribution of $[RA]_{out}$ in the morphogen domain and the higher magnification A-P distribution of $[RA]_{in}$ for the A-P range present in the tissue domain. The curves show the average morphogen level along the L-R axis. Cases with **(E)** or without **(F)** advections in $[RA]_{in}$ and $[FGF]_{signal}$.
